# Extracting host-specific developmental signatures from longitudinal microbiome data

**DOI:** 10.1101/2025.11.22.689760

**Authors:** Balázs Erdős, Christos Chatzis, Jonathan Thorsen, Jakob Stokholm, Age K. Smilde, Morten A. Rasmussen, Evrim Acar

## Abstract

Longitudinal microbiome studies provide critical insights into microbial community dynamics and their relation to host health. Tensor decompositions offer a powerful framework for the unsupervised analysis of such data, yielding interpretable low-dimensional temporal patterns. However, existing approaches based on the CANDECOMP/PARAFAC (CP) model assume common temporal dynamics for all subjects and therefore cannot capture subject-specific trajectories. To address this limitation, we introduce a novel analytical framework based on PARAFAC2 to explicitly model subject-specific variations, such as shifts and delays in temporal patterns. Through systematic comparisons on simulated and real-world datasets—including studies of infant gut maturation and dietary interventions—we demonstrate that PARAFAC2 outperforms CP in capturing subject-specific temporal trajectories, and enables the discovery of biologically relevant patterns that are overlooked by CP. Furthermore, we introduce replicability as a robust criterion for selecting the number of components and demonstrate that the extracted patterns are replicable.

**Author summary:** Longitudinal microbiome datasets are complex, consisting of repeated high-dimensional compositions that track changes in microbial abundance across individuals over time. While tensor decompositions are powerful tools for unraveling structure in these data, standard models like CANDECOMP/PARAFAC (CP) impose a critical limitation: they assume that temporal dynamics are identical across all individuals. This assumption often fails to capture the heterogeneity of biological processes, such as the varying pace of gut microbiome maturation or distinct individual responses to dietary changes. In this work, we introduce a novel analytical framework leveraging the PARAFAC2 model to overcome these constraints. By explicitly modeling subject-specific temporal variations, our PARAFAC2-based approach allows for the detection of individual time shifts and delays. We validated this framework using both simulation and real-world cohorts, demonstrating its ability to recover personalized trajectories that CP obscures. Additionally, we implemented a robust criterion to guide model selection, ensuring that the discovered patterns are replicable.

## Introduction

The human microbiome exhibits complex temporal dynamics that vary substantially both within and across individuals, making longitudinal analysis crucial for understanding its evolution through developmental stages (e.g., through childhood) or following interventions, as well as its link to host health. Advances in high-throughput sequencing now allow for dense temporal sampling, offering new opportunities to link microbial dynamics with host phenotypes. However, analyzing such data poses distinct computational challenges that call for specialized methods [1]. Traditional dimensionality reduction methods, such as principal coordinates analysis (PCoA) [2], operate at the sample level and fail to account for the inherent temporal structure and within-subject correlations present in longitudinal data.

To address these limitations, tensor decomposition-based approaches have recently emerged for the analysis of longitudinal microbiome data. Tensor decompositions, which are extensions of matrix factorizations to multi-way arrays (i.e., higher-order tensors), factorize multi-way data into a set of interpretable components, with applications across diverse fields such as chemometrics, neuroscience, and computational phenotyping [3–5]. In microbiome research, existing tools such as compositional tensor factorization (CTF) [6], parafac4microbiome [7], microTensor [8], and temporal tensor decomposition (TEMPTED) [9] have primarily relied on the CANDECOMP/PARAFAC (CP) decomposition [10, 11], which represents the data as a sum of components, each defined by taxa loadings (a microbial signature), subject loadings (its expression across individuals), and time loadings (its temporal progression).

However, a critical limitation of CP-based methods is the assumption of shared time loadings across all subjects. This assumption restricts these methods to modeling only population-level dynamics and prevents them from capturing subject-specific temporal variation that is commonly observed in microbiome studies [12]. This limitation is particularly relevant when individual temporal variation may reflect important biological information. For instance, in studies of infant gut microbiome maturation, subjects may follow the same underlying maturation process but at different paces, representing biological heterogeneity or environmental exposures [13]. In intervention studies involving antibiotics or dietary changes, subjects often exhibit divergent responses in the timing and magnitude of microbial adaptation (even within the same treatment group), underscoring the need for models that can capture these personalized dynamics [14, 15].

Unlike the CP model, PARAFAC2 [16] offers a more flexible alternative by relaxing the strict assumption of multilinearity when analyzing multi-way data by letting the factors in one mode change across tensor slices. This allows it to model shape changes, such as the shifts and delays that characterize individualized temporal trajectories. The power of this flexible approach has been well established in other domains. In chemometrics, for instance, PARAFAC2 is used to resolve and quantify chemical compounds in mixtures where chromatographic profiles exhibit retention time shifts between runs [17]. In neuroscience, it successfully identified changes in brain connectivity consistent with prior pathological knowledge of Alzheimer’s disease [18] and captured subject-specific temporal profiles from the analysis of multi-subject functional neuroimaging data [19]. It has been applied to electronic health records to construct personalized disease trajectories, capturing phenotypic evolution across clinical encounters [20]. More recently, in single-cell genomics, PARAFAC2 has provided a framework for integrating data from perturbational experiments, capturing how cellular responses vary across different conditions and treatments [21].

In this work, we adopt PARAFAC2 to analyse longitudinal microbiome data, allowing us to model subject-specific temporal trajectories. We evaluate and compare CP and PARAFAC2 on both simulated and real-world longitudinal microbiome datasets, including studies on infant gut maturation and dietary interventions. We show that PARAFAC2 more effectively captures subject-specific temporal dynamics, which in turn enables the discovery of biologically relevant patterns that are overlooked by the more restrictive CP model. Finally, we illustrate and discuss the use of replicability analyses in model selection to improve the robustness of the extracted patterns.

## 1 Materials and methods

### 1.1 CANDECOMP/PARAFAC (CP)

The CP model [10, 11, 22] approximates a higher-order tensor as the sum of rank-one tensors. An *R*-component CP model of a third-order tensor 𝒳*∈* ℝ^*I*×*J*×*K*^ is defined as 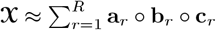, where *◦* denotes the vector outer product, **a**_*r*_, **b**_*r*_, and **c**_*r*_ correspond to the *r*-th column of factor matrices **A** *∈* ℝ^*I*×*R*^, **B** *∈* ℝ^*J*×*R*^, and **C** *∈* ℝ^*K*×*R*^, respectively. Using the CP model, the *k*th frontal slice of 𝒳, denoted **X**_*k*_ = 𝒳_:,:,*k*_, is approximated as:

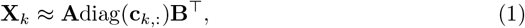

where diag(**c**_*k*,:_) is an *R*×*R* diagonal matrix with the *k*th row of **C** on the diagonal, and ^**⊤**^ denotes the matrix transpose. The CP model is unique up to permutation and scaling ambiguities under mild conditions [23], ensuring reliable interpretation of the resulting factors. More specifically, columns of the factor matrices can be permuted in the same way across modes (permutation ambiguity), and can be scaled as long as the component norm stays the same, e.g.,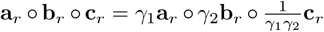 (scaling ambiguity). Under these ambiguities, the interpretation of the CP factors remains unchanged. For a *taxa* by *time* by *subjects* tensor 𝒳, each column of **A** (taxa loadings), i.e., **a**_*r*_, may reveal a microbial signature with the corresponding temporal profile captured in columns of **B** (time loadings), i.e., **b**_*r*_. Columns of **C** (subject loadings), i.e., **c**_*r*_, indicate the extent to which the compositional changes over time are present in each subject (Fig 1).

**Fig 1.**
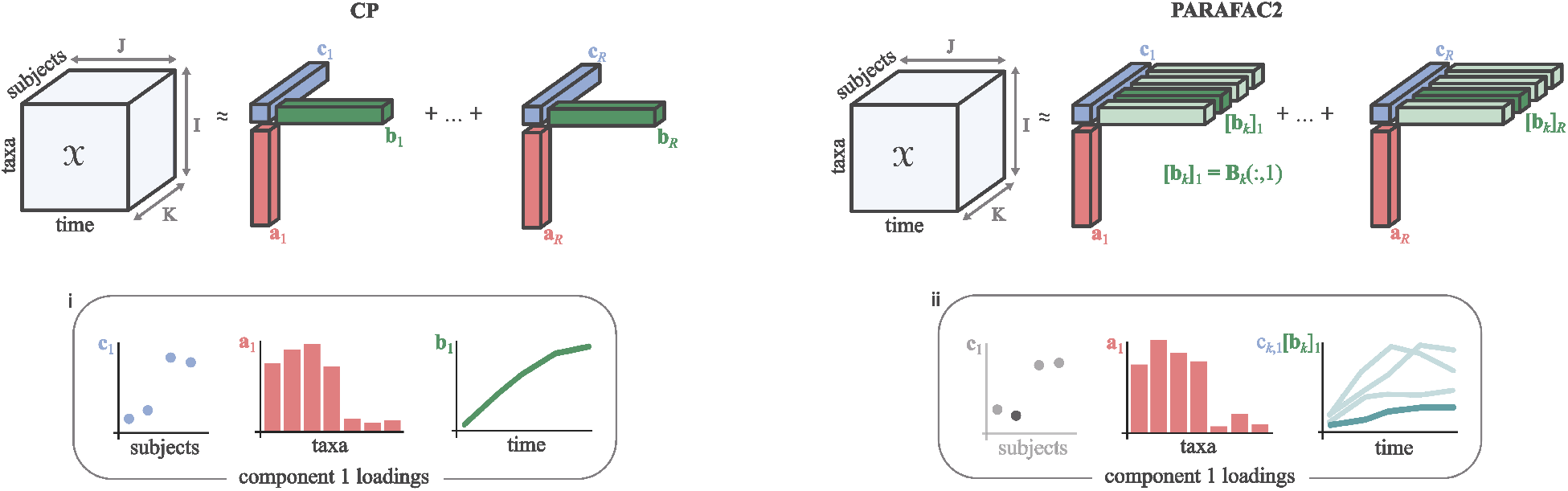
Illustration of an *R*-component CP (left) and PARAFAC2 (right) model of longitudinal microbiome data arranged as a third-order tensor 𝒳*∈* ℝ^*I*×*J*×*K*^ . PARAFAC2 decomposes 𝒳into taxa factors **A** = [**a**_1_, …, **a**_*R*_], subject factors **C** = [**c**_1_, …, **c**_*R*_], and host-specific time factors **B**_*k*_ = [**b**_*k*,1_, …, **b**_*k,R*_] ∀*k* ≤ *K*, where *K* is the number of subjects. PARAFAC2 allows the **B**_*k*_ factor matrices to vary across tensor slices under the constraint that their cross product is constant. This is more flexible than CP, which finds common time factors **B**_1_ = **B**_2_ = … = **B**_*K*_ = **B** = [**b**_1_, …, **b**_*R*_] describing all subjects. Component *r* comprises taxa loadings **a**_*r*_ characterizing a microbial signature, time loadings **b**_*r*_ in case of CP, and [**b**_*k*_]_*r*_ = **B**_*k*_(:, *r*) with PARAFAC2 that capture how the signature evolves over time, and subject loadings **c**_*r*_ that reflect the strength of the microbial signature over time in each individual. Example visualization of a component across its three modes (i), and in two modes, after deriving *scaled time loadings* by scaling the time loadings with their corresponding subject loading, i.e. *c*_*k,r*_[**b**_*k*_]_*r*_ ∀*k* ≤ *K* (ii). Scaled time loadings in case of CP are calculated as *c*_*k,r*_**b**_*r*_ ∀*k* ≤ *K*.

### 1.2 PARAFAC2

Unlike the CP model, the PARAFAC2 model [16] allows the patterns in **B**_*k*_ to vary across the third mode:

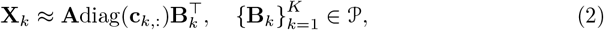

where 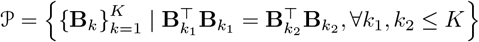 is the set defining the constant cross-product constraint of PARAFAC2, which ensures unique decomposition up to scaling and permutation ambiguities under certain conditions [24]. The added flexibility leads to the additional challenge of sign ambiguity in PARAFAC2. Reformulating Eq (2) as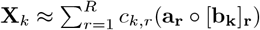, where *c*_*k,r*_ = **C**(*k, r*), **a**_*r*_ = **A**(:, *r*), and [**b**_*k*_]_*r*_ = **B**_*k*_(:, *r*), we observe that *c*_*k,r*_ may arbitrarily flip signs together with [**b**_*k*_]_*r*_ [16].

We resolve this ambiguity by imposing non-negativity on **C** [24]. Given a *taxa* by *time* by *subjects* tensor 𝒳, PARAFAC2 relaxes the strict assumption of the CP model by allowing temporal patterns **B**_*k*_ to vary across subjects *k*. In contrast, CP requires identical temporal patterns for all subjects, i.e, **B**_1_ = **B**_2_ = … = **B**_*K*_ = **B** (Fig 1). Therefore, the PARAFAC2 model may be better suited for datasets in which subjects exhibit heterogeneous temporal patterns.

Taxa, time, and subject loadings of the CP and PARAFAC2 models are visualized in Fig 1, either using all three modes (panel i), or in two modes (panel ii), by scaling the time loadings with their corresponding subject loading, referred to as scaled time loadings, i.e. *c*_*k,r*_[**b**_*k*_]_*r*_ ∀*k* ≤ *K*.

### 1.3 Model selection

Selecting the appropriate number of model components *R* is crucial for ensuring that the resulting model is both biologically meaningful and valid. Several approaches have been proposed to determine the number of components [25–27]; nevertheless, this task remains an active topic of research. In this study, we determine *R* based on the replicability and interpretability of the extracted components [27, 28]. Replicability refers to the consistency of identifying similar patterns across random subsamples of the dataset and extends the concept of split-half analysis [29]. We adopt a replicability-based model selection approach—by subsampling along the subject mode—to ensure that the extracted patterns are robust and representative of the underlying study population, as previously done to select the number of components for CP models [30]. Let 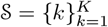 be the set of all slice indices in the subject mode of the complete data tensor. As an example, consider the subsampled tensors, 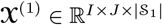 and 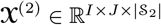, formed by stacking the slices {**X**_*k*_ | *k* ∈ 𝒮_1_} and {**X**_*k*_ |*k* ∈𝒮_2_}, respectively, where 𝒮_1_ ⊂ 𝒮and 𝒮_2_ ⊂ 𝒮are randomly selected, partially overlapping subsets of indices and |𝒮| is the cardinality of the set 𝒮. To quantify the similarity between factors extracted from different subsampled tensors (e.g. 𝒳^(1)^ and 𝒳^(2)^), we use the factor match score (FMS). For two CP models, e.g., CP models of 𝒳^(1)^ and 𝒳^(2)^, FMS over the taxa and time modes (FMS_AB_) is computed as:

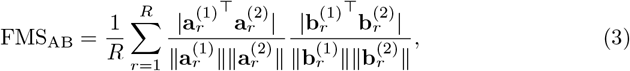

where ∥.∥ denotes the vector 2-norm, 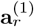 and 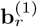 are the *r*-th columns of the taxa and time factor matrices from the CP model of 𝒳^(1)^ (first submodel), and 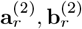, are the corresponding columns from the CP model of 𝒳^(2)^ (second submodel).

In the case of PARAFAC2, we compute FMS_A_ using the relevant terms (i.e. taxa mode loadings **a**_*r*_) of Eq 3. Since the time mode factors are specific to the set of subjects in the data tensor, they can only be compared using the loading vectors corresponding to the set of subjects common to any two submodels. To do this, we define maps such as *π*_1_ : 𝒮_1_ → {1, …, |𝒮_1_|} and *π*_2_ : 𝒮_2_ → {1, …, |𝒮_2_|} from a position in the complete tensor to its positional index in the submodel. Let 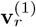 and 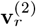 be the vectors of vertically concatenated scaled time loadings of the subjects shared between submodels one and two:

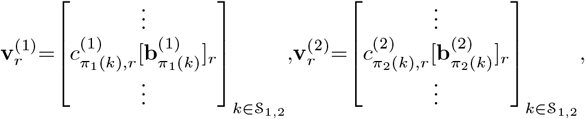

where 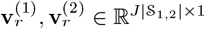 and 𝒮= 𝒮_1,2_ ∩ 𝒮_1_ . We then calculate FMS_C*B_ as

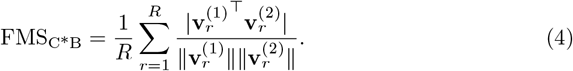

The factor match score is calculated over the scaled time loadings to avoid overemphasizing poorly estimated time loadings, which can arise when the corresponding subject loading is close to zero [31]. Because of the permutation ambiguity in CP and PARAFAC2 decompositions, we first align the order of components to the optimal permutation before computing FMS. The FMS ranges from 0 to 1, with higher values indicating greater similarity. Specifically, we assess the replicability of an *R*-component CP and PARAFAC2 model as follows.

1. Randomly split the dataset along the subjects dimension into *F* approximately equal parts, preserving class proportions if class labels are relevant.
2. For each part *f* = 1, …, *F* create a subsampled tensor by excluding the *f* -th part.
3. Fit an *R*-component submodel to each of the *F* subsets.
4. Compute factor similarity for all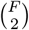 pairs of the *F* submodels, using FMS_AB_ for CP, and both FMS_A_ and FMS_C*B_ for PARAFAC2.
5. Repeat (1-4) ten times.

The number of subsets *F* depends on the characteristics of the dataset in use, such as the number of subjects *K* and the presence of relevant grouping labels. We consider an *R*-component model to be replicable if at least 90% of the pooled FMS_AB_ (or FMS_A_ and FMS_C*B_ in the case of PARAFAC2) values exceed 0.9. To maximize explanatory power, we select the largest number of components that still yield a replicable model. Beyond replicability, we adopt an exploratory approach to assess biological relevance [27].

### 1.4 Datasets

#### 1.4.1 Simulation study

In order to illustrate the use of PARAFAC2 to capture subject-specific temporal trends in longitudinal microbiome data, we constructed a simulated data tensor including variation in the temporal patterns across subjects. The data were generated in a way that does not fully conform to the structural assumptions of either CP or PARAFAC2, to better represent real-world scenarios where the true data-generating process is unknown. The simulated data tensor 𝒳∈ ℝ^*I*×*J*×*K*^ with *I* = 36 microbes, *J* = 21 time points, and *K* = 12 subjects was constructed according to the ground truth factors shown in Fig 2. Each of the three patterns, corresponding to the *R* = 3 components, describes the evolution of a microbial signature **a**_*r*_ over time according to the temporal profile [**b**_*k*_]_*r*_ with strength *c*_*k,r*_ in subject *k*. All three components are constructed to encode group-level differences between subsets of subjects through the subject loadings (**c**_*r*_). In addition, the second component includes subject-specific delays in the temporal profiles ([**b**_*k*_]_2_), through variation introduced to the time loadings across a subset of subjects. In particular, the first pattern is the bell-shaped trend in time of taxa 1-12 and is unique to the group of subjects {1,2,6,7,11,12} . The third pattern is the dual-peaking temporal trend of microbes 19-30 and is unique to the group of subjects {7,8,9,10,11,12} . The second pattern is the subject-specific saturation of microbes 10-21 that is unique to each subject within the group {4,5,6,7,8,9} . Microbes 31-36 have low feature loadings in all subjects at all time points. Of interest are the between-group differences, with only one group of subjects exhibiting the taxa signatures in each pattern, as well as the between-subject differences that are specific to a group of subjects in component 2. Simulated tensors were constructed both in the absence and presence of noise. To introduce noise, we constructed a noisy tensor as

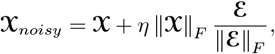

where 𝒳denotes the noiseless tensor, ℰ ∼ 𝒩 (0, 1) is a tensor of i.i.d. Gaussian noise, ∥.∥_*F*_ denotes the Frobenius norm, and the noise level was set to *η* = 0.25.

**Fig 2.**
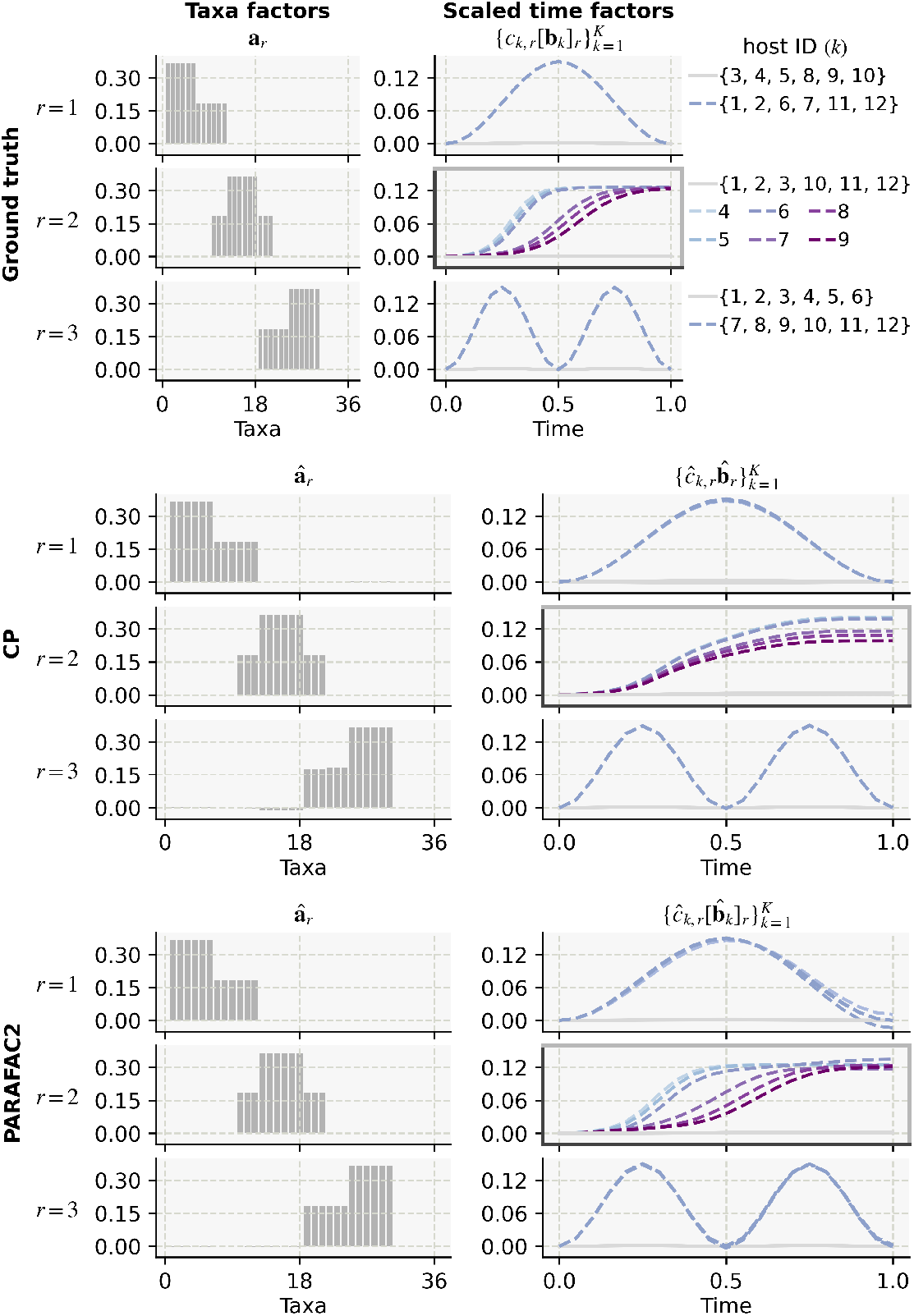
Ground truth factors underlying the simulated dataset and their recovery by CP and PARAFAC2. Top panel: ground truth factors including the taxa (**a**_*r*_), time ([**b**_*k*_]_*r*_), and subjects (**c**_*r*_) loadings visualized after scaling the time loadings by the corresponding subject loading (*c*_*k,r*_[**b**_*k*_]_*r*_). The microbial signature **a**_*r*_ is present in subject *k* over time according to their scaled time loadings *c*_*k,r*_[**b**_*k*_]_*r*_. Group differences characterized by a presence/absence of the microbial signature in the subject is indicated by dashed colored/continuous gray lines. Thick borders highlight the component 2 subpanels containing the patterns of interest. These comprise subject-specific time loadings *c*_*k*,2_[**b**_*k*_]_2_ for *k* ∈ {4, 5, 6, 7, 8, 9} as denoted by the respective line colors. Middle and bottom panels: factors recovered by CP and PARAFAC2, respectively.

#### 1.4.2 COPSAC_2010_ cohort

The Copenhagen Prospective Studies on Asthma in Childhood 2010 (COPSAC_2010_) is a population-based birth cohort of 700 children recruited in pregnancy and followed prospectively with deep clinical phenotyping [32]. The study was conducted in accordance with the guiding principles of the Declaration of Helsinki and was approved by the Local Ethics Committee (H-B-2008-093), and the Danish Data Protection Agency (2015-41-3696). Both parents gave written informed consent before enrollment. The data used here comprises fecal samples collected at 1 week, 1 month, 1 year, 4 years, and 6 years after birth and analyzed via 16S rRNA gene amplicon sequencing.

Details of the laboratory workflow, sequencing and data processing have been described previously [33, 34]. Additional covariates include mode of delivery (vaginal delivery/Cesarean section) and maternal intrapartum antibiotics (yes/no). Children with missing samples were excluded from the analysis. Species not detected in ≥ 10% of subjects in at least one time point were filtered out. To account for the compositional nature of the data, the centered log-ratio (CLR) transformation was applied, using a pseudo-count of 0.5 [35]. After pre-processing and data transformation, the data were arranged as a third-order tensor of size 364× 5× 267 with modes taxa, time, and subjects, respectively. The data tensor contained no missing entries.

#### 1.4.3 FARMM study

The Food and Resulting Microbial Metabolites (FARMM) study collected daily fecal samples over a 15-day period from 30 adult participants, evenly distributed across three diet groups: vegan, omnivore, and exclusive enteral nutrition (EEN) [15]. The study comprised three distinct phases: the dietary phase (days 1–5), the gut microbiota purge phase (days 6–8), and the recovery phase (days 9–15). All participants received antibiotic and polyethylene glycol (Abx/PEG) treatment during days 6 to 8. For a detailed description of the study design, sample collection, and sequencing procedures, we refer to the original publication [15]. Preprocessed shotgun metagenomics-based relative abundance data were obtained from [8]. Taxa detected in fewer than five samples with a relative abundance of at least 10^−5^ were filtered out. Additionally, time point zero was removed because no subjects in the vegan group had samples at that time point. Subsequently, the CLR transformation was applied after adding a pseudo-count of 0.5. The data were arranged as a third-order tensor of size 343 × 15 × 30 with modes taxa, time, and subjects, respectively. The data tensor contained 11.8% missing entries.

### 1.5 Experimental setup

#### 1.5.1 Implementation details

Before applying the tensor decompositions, data tensors were divided by their Frobenius norm. For fitting the CP models, we used an alternating least squares-based algorithm [36] implemented in Tensorly (v0.9.0) [37]. The PARAFAC2 models were fit using alternating optimization with the alternating direction method of multipliers (AO-ADMM) algorithm implemented in MatCoupLy (v0.1.6) [31, 38]. In experiments with missing data, we used an expectation maximization (EM)-based approach with both CP and PARAFAC2 [37, 39]. We applied the following constraints and regularization terms:

- **Simulation study:** Non-negativity in the subjects mode (i.e., on factor matrix **C**) in PARAFAC2.
- **COPSAC_2010_ data:** Ridge penalty in all modes in CP. Non-negativity in the subjects mode and ridge penalty in all modes in PARAFAC2.
- **FARMM data:** Ridge penalty in all modes in CP. Non-negativity in the subjects mode, ridge penalty in the taxa and subjects modes (i.e., **A, C**), and graph Laplacian penalty in the time mode (i.e.,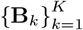) in PARAFAC2.

The ridge penalty was added to aid convergence [40], using a penalty parameter set to 10^−3^. Non-negativity was imposed to resolve the sign ambiguity of PARAFAC2. Due to missing samples, the FARMM dataset contains completely missing mode-1 fibers (i.e., all taxon relative abundances **X**_*k*_(:, *j*) for subject *k* at time point *j* are missing). PARAFAC2 struggles with such scenarios, as **B**_*k*_(*j*, :) can be chosen arbitrarily, provided it conforms to the imposed constraints. To improve the robustness of the recovered **B**_*k*_ factor matrices, we used a smoothness constraint in the form of a graph Laplacian penalty (with penalty parameter 10^−3^) on columns of the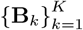 factor matrices to penalize large differences in neighboring time points [31]. To avoid convergence to local minima, each model was fit using multiple random initializations. After confirming the uniqueness of the solution, the solution with the best fit was selected for further analysis. Finally, to address the scaling ambiguity of CP and PARAFAC2, we scaled the columns of factor matrices to unit norm.

#### 1.5.2 Performance metrics

To evaluate how well CP or PARAFAC2 models the data, we calculated the model fit as follows:

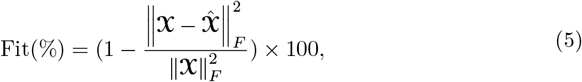

where 𝒳is the data tensor,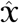 is its reconstruction based on the model. A model fit of 100% indicates that the model perfectly reconstructs the data, while a fit below 100% implies that some variation in the data remains unexplained.

For simulations where the ground truth factors were known, we used FMS to assess how accurately the ground truth factors were recovered. Here, we used FMS across all modes, defined as:

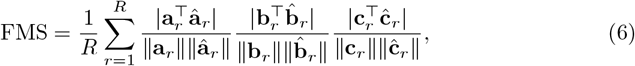

where **a**_*r*_, **b**_*r*_, and **c**_*r*_ are the *r*-th columns of the ground truth factor matrices, and **â**_*r*_,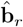, and **ĉ**_*r*_ are the corresponding model estimates. In the case of PARAFAC2, **b**_*r*_ and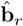 refer to vectors produced by vertically concatenating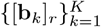 and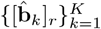, respectively.

## 2 Results

### 2.1 PARAFAC2 recovers subject-specific temporal trends

To demonstrate the utility of PARAFAC2 in microbiome data analysis, we compared its performance with that of CP in recovering the underlying true factors from a simulated dataset specifically designed to reflect subject-specific variation in the temporal patterns. The ground truth as well as the recovered factors are shown in Fig 2.

While both models achieved high overall fit (CP: 98.8%, PARAFAC2: 99.9%) as well as factor match scores (CP: 0.994, PARAFAC2: 0.997) and successfully recovered the common taxa factors (**a**_1_, **a**_2_, **a**_3_) and temporal dynamics in components 1 and 3, they differed substantially in handling subject-specific variation. CP did not recover the subject-specific time loadings of comp. 2 due to its assumption of shared time loadings between subjects (Fig 2, middle panel, highlighted with solid borders); the recovered profiles misrepresented the trajectories as identical across subjects, with only scale differences due to subject loadings *c*_*k*,2_. PARAFAC2, however, successfully captured the between-subject variability in comp.

2. In particular, it faithfully recovered the temporal patterns of subjects 4-9 (Fig 2, bottom panel, highlighted). While CP forced a shared temporal profile, PARAFAC2 accurately resolved the saturation timing differences, achieving a mean absolute error in 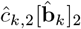 of 0.0013 versus 0.0068 for CP. This demonstrates the superior performance of PARAFAC2 in capturing subject-specific dynamics. Results were similar, with PARAFAC2 outperforming CP at 25% noise (S2 Fig).

### 2.2 Subject-specific maturation patterns of the infant gut microbiome in the COPSAC_2010_ cohort

To characterize infant gut microbiome maturation while accounting for subject-specific differences in the temporal evolution of dominant microbial signatures, we applied the PARAFAC2 decomposition to 16S rRNA longitudinal gut microbiome data from 267 children of the COPSAC_2010_ cohort. To highlight the distinctions in the extracted patterns, we also compared the results with those obtained using the CP decomposition. We focus on the model components capturing microbial variation in early life and related to mode of delivery and maternal intrapartum antibiotic exposure to illustrate the approaches. For downstream analysis and visualization, subjects were grouped into exposure groups according to mode of delivery and maternal intrapartum antibiotics: (i) vaginal delivery without intrapartum antibiotic prophylaxis (VD-no IAP), (ii) vaginal delivery with intrapartum antibiotic prophylaxis (VD-IAP), and (iii) Cesarean section (with intrapartum antibiotic prophylaxis; CS).

We selected a 5-component PARAFAC2, and a 7-component CP model based on our model selection approach, focusing on replicability and interpretation of the components (S3 Fig). The replicability analysis was conducted using *F* =10 subsets, generated through sampling stratified by delivery mode to account for its known impact on gut microbiome development [41, 42]. For PARAFAC2, the dimensions of the dataset limited the maximum number of extractable components to five, in line with the model’s uniqueness conditions [24]. Notably, PARAFAC2 explained slightly more variance (50.0%) with fewer components compared to CP (49.8%), reflecting its ability to capture temporal misalignment more efficiently. Both models identified components that capture early-life patterns—specifically, variation in samples collected at 1 week, 1 month, and 1 year of age (PARAFAC2: comp. 3 & 5, Fig 3; CP: comp. 2 & 7, S6 Fig). The complete models, including all components, are visualized in S4 Fig and S5 Fig.

**Fig 3.**
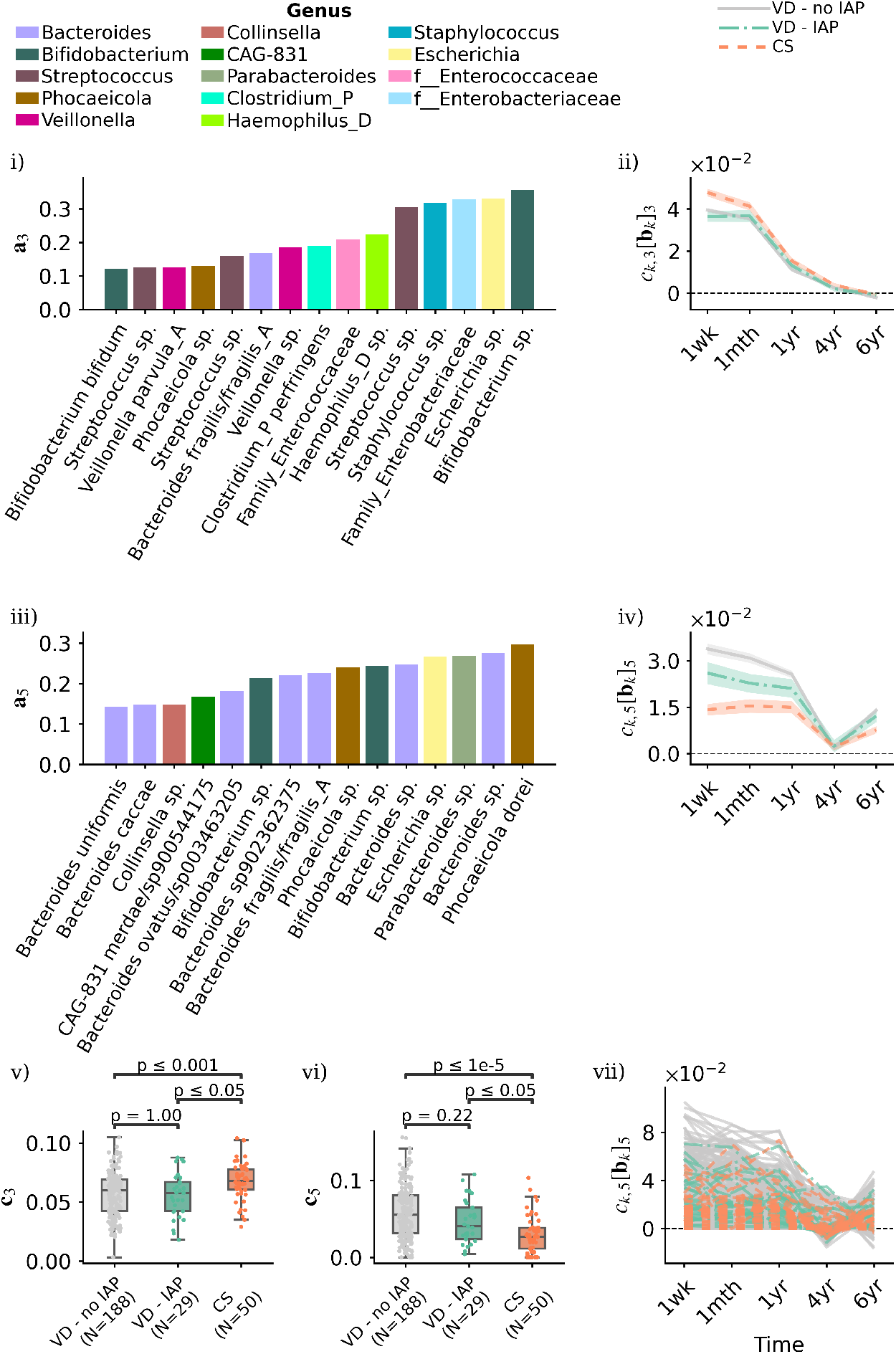
PARAFAC2 uncovers subject-specific temporal patterns of gut microbial signatures in early life, related to delivery mode and maternal intrapartum antibiotics in the COPSAC_2010_ cohort. Components 3 (i–ii) and 5 (iii–iv) capture patterns in gut microbiome composition during the first year of life. Panels (i, iii) show taxa loadings with the 15 ASVs of largest absolute weight in the component, colored by genus. Scaled time loadings (ii, iv) at time points were averaged per exposure group. Shaded area indicate standard errors of group means. Panels (v, vi) present subject loadings in components 3 and 5 compared across exposure groups using the Mann–Whitney U test with Bonferroni correction. Panel (vii) illustrates subject-specific scaled time loadings of component 5, highlighting heterogeneity in the trajectories.

The early-life representations of the PARAFAC2 and CP models were broadly consistent in terms of the uncovered microbial signatures (Fig 3, panels i, iii; S6 Fig, panels i, iii). The top 15 taxa with the largest absolute weights in each of the early-life-related components (PARAFAC2 comp. 3 vs. CP comp. 2, and PARAFAC2 comp. 5 vs. CP comp. 7) showed an 87% and 80% overlap between the models at the ASV level. When considering the union of top 15 ASVs across the components, the overlap between models is 89%, highlighting the consistency in the extracted microbial signatures. However, while the taxa profiles were similar, the models differed in their ability to resolve group-level and subject-specific temporal dynamics (Fig 3, panels ii, iv; S6 Fig, panels ii, iv).

Both PARAFAC2 and CP recovered a component representative of a general early-life microbial signature (PARAFAC2 comp. 3 and CP comp. 2), characterized by typical early colonizers such as *Escherichia* species and members of the Enterobacteriaceae family (Fig 3, panel i; S6 Fig, panel i) [43]. In both models, the scaled time loadings show that the prevalence of this signature decreases gradually over time (Fig 3, panel ii; S6 Fig, panel ii). However, the added flexibility of PARAFAC2 revealed more differential dynamics across exposure groups that CP failed to resolve. While CP subject loadings indicated a difference between VD-no IAP and VD-IAP (S6 Fig, panel v), PARAFAC2 distinguished VD (both without and with IAP) from CS (Fig 3, panel v). Furthermore, the PARAFAC2 scaled time loadings *c*_*k*,3_[**b**_*k*_]_3_ captured a temporal interaction: the microbial signature was more dominant in CS compared to VD specifically at 1 week (Fig 3, panel ii), a detail obscured in the CP model.

The second early-life component (PARAFAC2 comp. 5 and CP comp. 7) captured a microbial signature with higher abundances of *Bacteroides* species in both models, accompanied in CP by a simultaneously decreased abundance of taxa such as *Clostridium perfringens* (Fig 3, panel ii; S6 Fig, panel ii). The subject scores for these signatures differed significantly by mode of delivery in both models (Fig 3, panel vi; S6 Fig, panel vi), consistent with known effects of Cesarean section [44, 45]. The CP model suggests that this signature diminishes over time in all groups (S6 Fig, panel iv), implying a uniform replacement of early colonizers [46, 47]. In contrast, the PARAFAC2 scaled time loadings show that the signature remains relatively stable in CS–born infants (characterized by consistently lower initial abundance), whereas in vaginally delivered infants, the initially high abundance decreases over time (Fig 3, panel iv).

This suggests that the uniform trajectory in CP may be an artifact of the model’s rigidity. Crucially, by examining subject-specific rather than group-mean profiles, PARAFAC2 unmasked substantial heterogeneity in gut microbiome maturation (Fig 3, panel vii; vs S6 Fig, panel vii). Unlike CP, which forces a shared temporal profile, PARAFAC2 successfully captured individual temporal shifts and delays, providing a more accurate representation of the subject-specific pace of microbiome development.

The microbial signatures of the early-life components captured by both models were relatively sparse (i.e., taxa loadings were characterized by few taxa), whereas the later-life components reflected taxonomically more diverse signatures (S4 Fig and S5 Fig), reflecting typical childhood gut microbiome development [43, 48].

### 2.3 Heterogeneous gut microbiome adaptation to dietary intervention in the FARMM study

We applied PARAFAC2 and CP to shotgun metagenomics-based longitudinal gut microbiome data from adults in the FARMM study to explore the effects of dietary and antibiotic/polyethylene glycol intervention (Abx/PEG) [15]. The replicability-based model selection approach resulted in a 3-component PARAFAC2 and a 3-component CP model (S7 Fig). Due to the smaller sample size (*N* = 10 in each of 3 study groups), the replicability analysis was conducted with *F* = 5 subsets, with stratified sampling for study group. The PARAFAC2 model explained a larger proportion of the variance in the data (51.0%) compared to CP (48.3%). Both models identified two components related to the dietary phase of the study (days 1-5) and one component related to the recovery phase (days 9-15) following the Abx/PEG perturbation (S8 Fig and S9 Fig).

The overall patterns of microbial features suggested a general concordance between the models across phases, as reflected in the substantial overlap of top-ranked species across components. In the first dietary phase–related component (comp. 1, both models; Fig 4), 87% of the top 15 species with the largest absolute weights overlapped between models. Overlap was complete (100%) for the recovery phase–related components (S10 Fig panels i, iv), with species typically associated with exposure to antibiotics such as *Veillonella* and *Klebsiella* being identified. The second dietary phase–related component showed greater divergence, with 60% overlap between the two models (S11 Fig). Here, the microbial signature in both models indicated an increased relative abundance of *Clostridium* species, while CP attributed larger absolute weights to the simultaneously decreased relative abundance of species, such as *Eubacterium eligens* and *Bifidobacterium adolescentis* (S11 Fig, panels i, iv). These patterns reflected the difference in microbial composition between the EEN group compared to the other diet groups which were consistent between PARAFAC2 and CP (S11 Fig, panels iii, vi).

**Fig 4.**
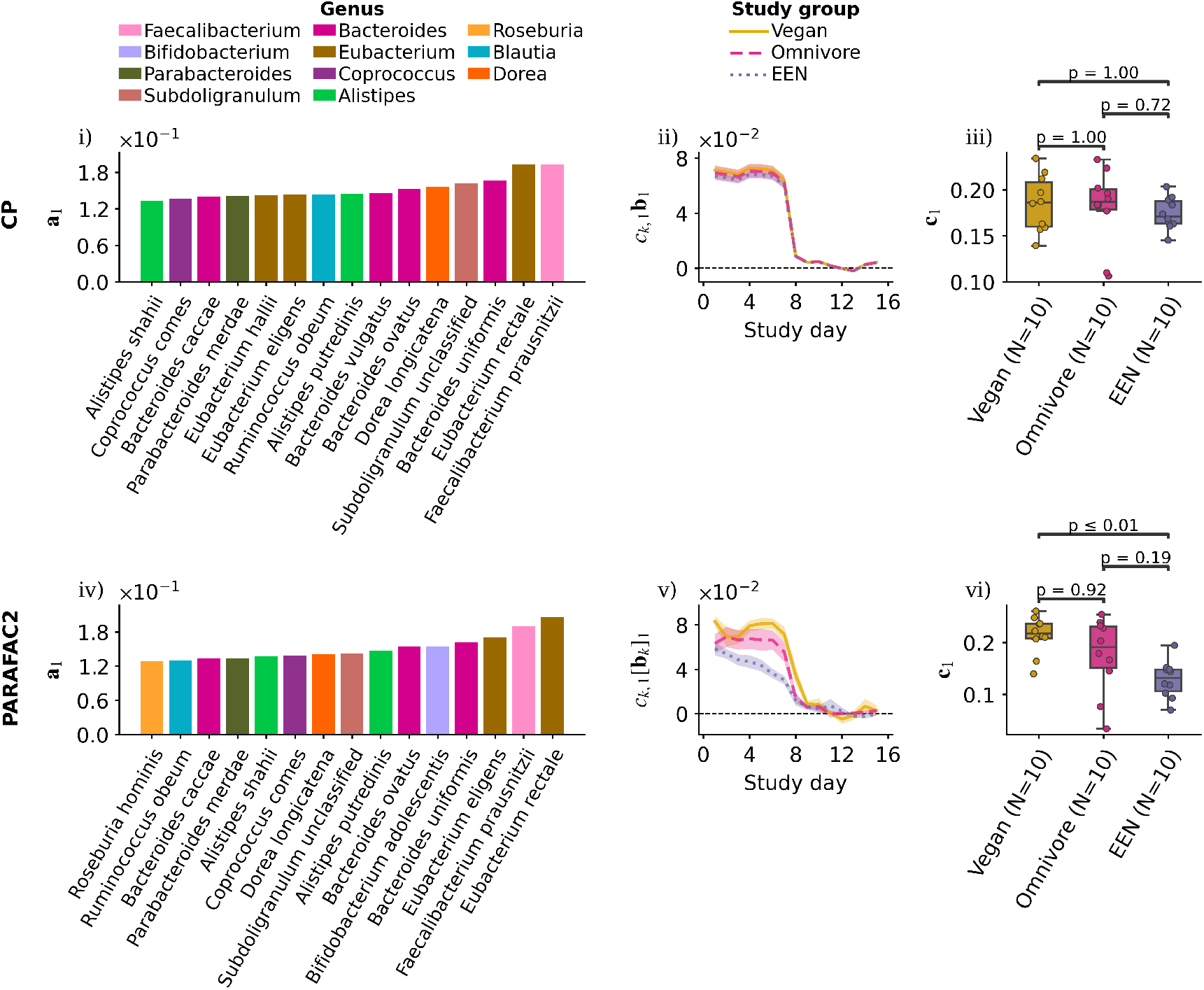
PARAFAC2 reveals distinct temporal patterns of gut microbiome response to dietary and antibiotic interventions in the FARMM study. Shown are component 1 of the selected CP (panels i–iii) and PARAFAC2 (panels iv–vi) models, associated with the dietary (days 1–5) and antibiotics phases (days 6–8) of the trial. Panels (i, iv) display the 15 species with the largest absolute weights in each component, representing the extracted microbial signatures. Scaled time loadings (ii, v) at time points were averaged per study group. Shaded area indicate standard errors of group means. Panels (iii, vi) present subject loadings compared across exposure groups using the Mann–Whitney U test with Bonferroni correction.

Both models identified broadly similar temporal trajectories of the microbial signatures, as reflected in the scaled time loadings; however, PARAFAC2 revealed subject-specific dynamics that were not apparent in the CP results. PARAFAC2 uncovered distinct microbial response to the exclusive enteral nutrition (EEN) diet compared to the vegan and omnivore diets in component 1, which was missed by CP (Fig 4, panel v). In particular, PARAFAC2 resolved a decrease in the abundance of the microbial signature during the dietary phase for the EEN group. This dynamic was not observed in the other groups. The improved resolution is further reflected in the corresponding subject loadings, where PARAFAC2 successfully distinguished between the EEN and vegan groups (Fig 4, panel vi). These findings align with those reported by Tanes et al. (2021) [15], highlighting the capability of PARAFAC2 to capture complex, individualized temporal patterns. Furthermore, while the CP model oversimplified the recovery phase as a constant difference across time, PARAFAC2 accurately modeled the evolving divergence between the EEN group and others (S10 Fig, panels ii, v).

### 2.4 Replicability of subject-specific factors

Beyond the global replicability (FMS_C*B_) used in model selection, we also assessed the uncertainty of subject-specific temporal patterns. To do this, we first identified, for each subject, the specific submodels from the replicability analysis in which they were included (89/100 and 39/50 submodels per subject for COPSAC_2010_ and FARMM, respectively). We then calculated the standard deviation of their scaled time loadings at each time point across the submodels and averaged the standard deviations to use as an indicator of the variance in their inferred profiles. The average standard deviations were generally an order of magnitude smaller than the scaled time loadings, indicating good agreement across submodels and robust recovery of individual temporal patterns (Fig 5, panels i, iii).

**Fig 5.**
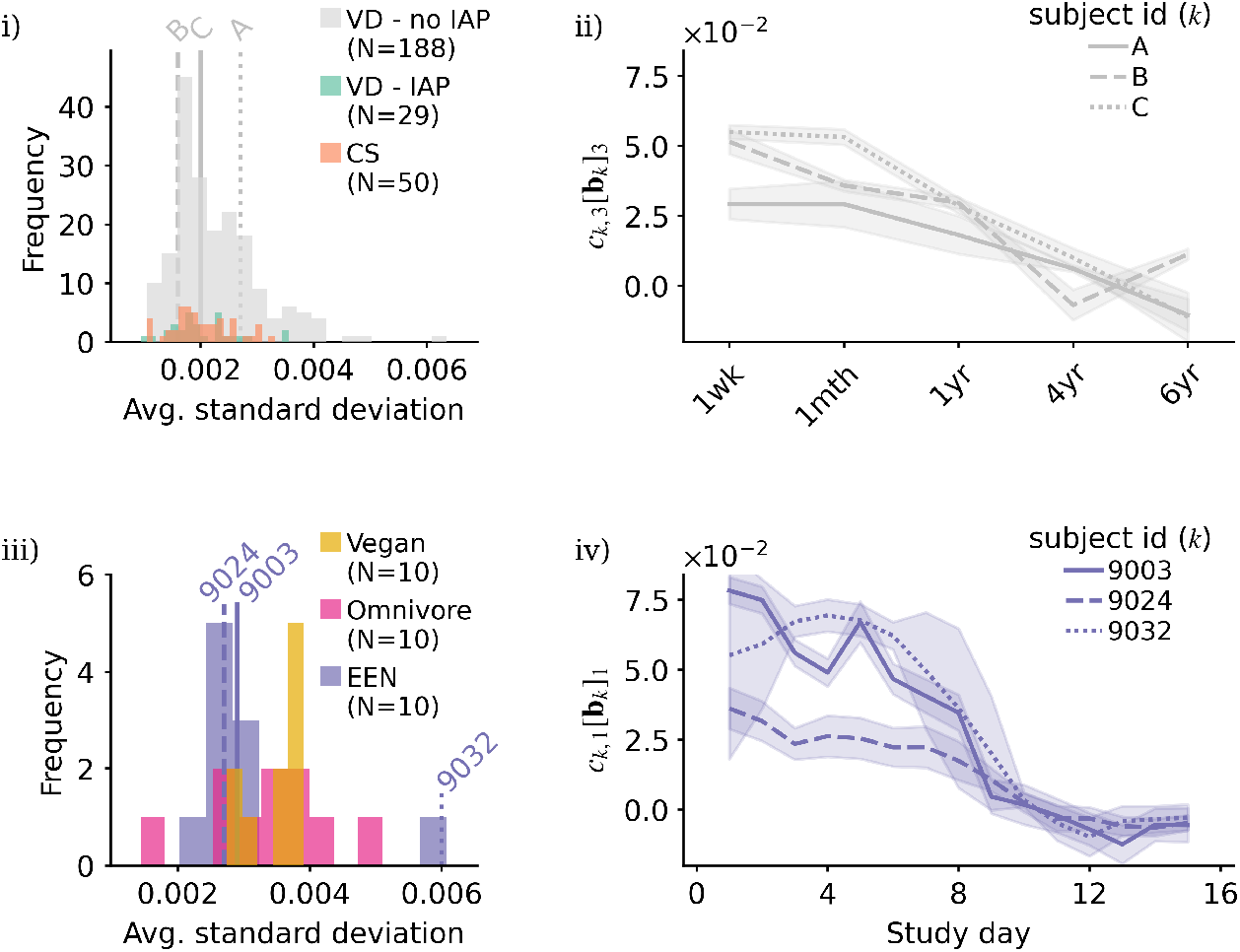
Replicability of PARAFAC2-based subject-specific scaled time loadings in the COPSAC_2010_ (top) and FARMM (bottom) datasets. Panels (i, iii) show histograms of the average standard deviation, calculated for each subject by first finding the standard deviation of the scaled time loadings at each time point across submodels, then taking the average. Panels (ii, iv) show the mean (lines) ±2 standard deviations (shaded area) of scaled time loadings at each time point across submodels for three representative subjects from the VD - no IAP group in the COPSAC_2010_ cohort (panel ii) and the EEN group in the FARMM (panel iv) study.

To illustrate the spectrum of uncertainty, we visualized the scaled time loadings from all relevant submodels for a selection of representative subjects from both cohorts (Fig 5). Overall, the profiles showed good agreement across submodels, confirming their replicability at the subject level, even within the same exposure group. For example, panel (ii) shows the mean (±2 standard deviations) for three subjects (A, B, C) in the VD - no IAP exposure group from the COPSAC_2010_ dataset, all of which show high consistency (subject-specific average FMS_C*B_ ≥ 0.99). Similarly, panel (iv) shows robust profiles for subjects 9003 and 9024 of the EEN study group from the FARMM dataset, with subject-specific average FMS_C*B_ ≥ 0.99. The comparatively larger uncertainty observed for subject 9032 (subject-specific average FMS_C*B_ = 0.97) reflects the underlying missingness pattern of the data, as this subject was missing samples on study days 1-2 and 7-9 (see Discussion). Results displaying the complete set of model components are shown in S12 Fig and S13 Fig, for the COPSAC_2010_ and FARMM studies, respectively.

## 3 Discussion

In this study, we introduced the PARAFAC2 model as a powerful tool for the unsupervised analysis of longitudinal microbiome data, specifically to capture subject-specific temporal trajectories. Furthermore, to ensure the robustness of the uncovered patterns, we proposed a replicability-based framework to determine the number of components, ensuring the biological relevance and stability of the extracted signatures.

Results from the simulated data expose a critical limitation of CP-based tensor decompositions for longitudinal microbiome data analysis: the inability to capture subject-specific microbial dynamics (such as accelerations or delays), phenomena frequently observed in real populations [49, 50]. This limitation was explicitly demonstrated by the peak timing of the microbial signature in component 2. While the ground truth exhibited a broad temporal shift, peaking early at *t* ≈ 0.4 for subject 4 and late at *t* ≈ 0.9 for subject 9, PARAFAC2 precisely resolved these distinct timings. In contrast, the CP model masked this heterogeneity, rigidly forcing a shared temporal profile that peaked at an intermediate *t* ≈ 0.75 for all subjects. Consequently, CP compensated for this timing misalignment by introducing variability in peak magnitude not supported by the data. In real-world contexts, where shifts in saturation timing reflect distinct environmental exposures or host states, accurate recovery of these individual trajectories is essential.

While PARAFAC2 outperformed CP in terms of fit and FMS, it is important to note that perfect scores are not expected, as the simulated data did not fully conform to the structural assumptions of either model. Since the true data-generating process is often unknown, the added flexibility of PARAFAC2 may make it the more suitable modeling choice when individual variation in temporal profiles is expected or of biological interest.

In real data, similarly to the simulation, PARAFAC2 outperformed CP in terms of fit, and the two models revealed broadly consistent microbial signatures. However, the scaled time loadings of PARAFAC2 provided a more fine-scale temporal evolution of these signatures at the individual level, as well as additional group differences not found by CP. The subject-specific PARAFAC2 patterns revealed heterogeneity among Cesarean-section-born infants, highlighting that some show an earlier change toward a composition resembling that of vaginally born infants. Identifying these subject-specific maturation trajectories is important, as they may reflect differences in postnatal exposures, including environmental conditions or feeding practices that help to compensate for the early-life perturbations [48, 51]. In the FARMM study, PARAFAC2 resolved a compositional change in the EEN group, characterized by decreasing relative abundances of species such as *Faecalibacterium prausnitzii, Parabacteroides merdae*, and *Bacteroides ovatus* during the dietary phase—dynamics consistent with [15], which were obscured by the CP model. The assumptions of CP require that all subjects follow the same temporal trajectory, allowing only for differences in the magnitude of how strongly each individual expresses the microbial signature according to the trajectory. Because of this assumption, inter-individual differences in the timing or progression of microbial changes could not be captured using CP, hindering the model’s ability to represent subject-specific dynamics and overlooking biologically meaningful patterns. Nevertheless, the increased complexity of PARAFAC2 may not always be necessary. For datasets or research questions where the primary goal is to characterize dominant, shared temporal trends rather than individual heterogeneity, the simpler CP model remains a robust and effective choice.

A limitation of PARAFAC2 lies in the constant cross-product constraint it imposes on the **B**_*k*_ factor matrices. Although the constraint is essential for obtaining a unique solution, which is required for interpretability, the assumption that loading vectors of **B**_*k*_ should have constant angles for all *k* may impose biologically unwarranted structure [52], potentially limiting the ability of the model to fully capture the diversity of subject-specific temporal dynamics. Additionally, the number of components PARAFAC2 can extract is limited by its uniqueness conditions. In particular, for datasets with only a few time points, a CP-based approach may be a better choice, as it may recover more microbial signatures. Nevertheless, in the analysis of functional magnetic resonance imaging (fMRI), PARAFAC2 has emerged as a robust compromise between models of differing flexibility, effectively capturing connectivity and spatial components while accounting for subject variability [53]. In addition, the model has been shown to perform well even when the PARAFAC2 constraint is only approximately satisfied, as demonstrated in our results on simulated data and in previous studies [31, 54].

An open challenge when fitting PARAFAC2 is the presence of completely missing measurements at a time point for a subject (i.e., a missing mode-1 fiber, such as in the case of the FARMM dataset), which makes the corresponding entries of the time loadings arbitrary as long as the imposed constraints are satisfied. To overcome this indeterminacy and improve the robustness of our results, we employed a smoothness constraint on the temporal mode. Nevertheless, in cases where such missingness is extensive (e.g., 9032, panel iv, Fig 5), this approach may be inadequate, highlighting the need for alternative strategies for the handling of missing mode-1 fibers.

Validating the subject-specific patterns of PARAFAC2 is challenging, owing to its added flexibility. In the absence of biological replicates, it is not possible to directly assess whether the subject-specific trajectories reflect true biological variation or are partly driven by model assumptions and data idiosyncrasies. To account for this, we focused on group-level trends, aggregating subject-specific loadings within predefined exposure groups (e.g., by delivery mode and diet group). While this approach captures consistent between-group differences, it may mask individual-level heterogeneity. Therefore, to more directly assess replicability at the subject level, we inspected the heterogeneity of subject-specific scaled time loadings between the submodels fitted during replicability analysis. By leveraging the partial overlap of subjects between submodels, we could compare subject-specific trajectories across submodels, which confirmed their replicability and supported our model selection approach. Nevertheless, parameters of the resampling, such as the number of subsamples, depend on dataset characteristics. Finally, replicate samples, when available, can aid in both determining the number of model components and assessing the biological validity of the discovered patterns. This approach has been demonstrated in metatranscriptomic analyses of microbial communities using CP [55], and the study of functional neuroimaging data via PARAFAC2 [19].

In summary, PARAFAC2 presents a flexible and interpretable approach to analyze longitudinal microbiome data while explicitly modeling subject-specific temporal trajectories. Accommodating timing differences, shifts and delays, PARAFAC2 enables a more accurate characterization of temporal changes. The integration of replicability as a criterion for model selection enhances the robustness of the extracted patterns.

Together, these features advance the methodological toolkit for studying microbiome dynamics and open avenues for future applications across diverse longitudinal omics datasets.

## Supporting information

**S1 Fig. Schematic of assessing the replicability of a CP or PARAFAC2 model with** *R* **components used in model selection**.

**S2 Fig. Ground truth factors and their recovery by CP and PARAFAC2**. Noise was added to the tensor as 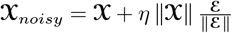 with ℰ ∼ 𝒩 (0, 1) and *η* = 0.25 where 𝒳is the simulated tensor. Top panel: ground truth factors including the taxa (**a**_*r*_), time ([**b**_*k*_]_*r*_), and subjects (**c**_*r*_) loadings with the time loadings scaled by the respective subject loading (*c*_*k,r*_[**b**_*k*_]_*r*_). The microbial signature **a**_*r*_ is present in subject *k* over time according to their scaled time loadings *c*_*k,r*_[**b**_*k*_]_*r*_. The presence/absence of the signature in the subject is indicated by blue dashed/gray continuous line. The patterns in component 2 contain subject-specific time loadings *c*_*k*,2_[**b**_*k*_]_2_ for *k* ∈{4, 5, 6, 7, 8, 9} as highlighted by the respective line colors. Middle and bottom panels: factors recovered by CP and PARAFAC2, respectively.

**S3 Fig. Selection of the number of components for CP and PARAFAC2 models of COPSAC**_**2010**_ **data**. Model fit (top row) and replicability (bottom row) in CP and PARAFAC2 models with increasing number of components (*R*). Box plots represent the median and interquartile range, with whiskers extending to the 10th and 90th percentiles. FMS: factor match score. While a 9-component CP model also satisfied our replicability criteria, the components related to early life and our exposures of interest were similar between the models, therefore we selected the 7-component CP model for parsimony.

**S4 Fig. Five-component PARAFAC2 model of the COPSAC**_**2010**_ **data**. Each row shows a component comprising taxa loadings (**a**_*r*_) and the time loadings scaled by their corresponding subjects loadings (*c*_*k,r*_[**b**_*k*_]_*r*_) for the *k*th subject and *r*th component.

**S5 Fig. Seven-component CP model of the COPSAC**_**2010**_ **data**. Each row shows the subject loadings (**c**_*r*_), taxa loadings (**a**_*r*_), and time loadings (**b**_*r*_) for the *r*th component.

**S6 Fig. Early life-related components of the selected CP model of the COPSAC**_**2010**_ **cohort**. Components 2 (i–ii) and 7 (iii–iv) capture shifts in gut microbiome composition during the first year of life. Panels (i, iii) show taxa loadings with the 15 ASVs of largest absolute weight in each component, colored by genus.

Scaled time loadings (ii, iv) at time points were averaged per exposure group. Shaded area indicate standard errors of group means. Panels (v, vi) present subject loadings in components 2 and 7 compared across exposure groups using the Mann–Whitney U test with Bonferroni correction. Panel (vii) illustrates subject-specific scaled time loadings of component 7.

**S7 Fig. Selection of the number of components for CP and PARAFAC2 models of FARMM study data**. Model fit (top row) and replicability (bottom row) in CP and PARAFAC2 models with increasing number of components (*R*). Box plots represent the median and interquartile range, with whiskers extending to the 10th and 90th percentiles. FMS: factor match score.

**S8 Fig. Three-component PARAFAC2 model of the FARMM study data**. Each row displays the taxa loadings (**a**_*r*_) together with the time loadings scaled by their corresponding subject loadings (*c*_*k,r*_[**b**_*k*_]_*r*_) for the *k*th subject and *r*th component.

**S9 Fig. Three-component CP model of the FARMM study data**. Each row displays the subject loadings (**c**_*r*_), taxa loadings (**a**_*r*_), and time loadings (**b**_*r*_) for the *r*th component.

**S10 Fig. Component 2 of the selected CP (panels i–iii) and PARAFAC2 (panels iv–vi) models of the FARMM dataset, associated with the recovery phase (days 9–15) of the trial**. Panels (i, iv) display the 15 species with the largest absolute weights in each component, representing the microbial signatures extracted by the models. Scaled time loadings (ii, v) at time points were averaged per study group. Shaded area indicates standard errors of group means. Panels (iii, vi) present subject loadings compared across exposure groups using the Mann–Whitney U test with Bonferroni correction.

**S11 Fig. Component 3 of the selected CP (panels i–iii) and PARAFAC2 (panels iv–vi) models of the FARMM dataset, associated with the dietary phase (days 1–5) and the antibiotics phase (days 6–8) of the trial**. Panels (i, iv) display the 15 species with the largest absolute weights in each component, representing the microbial signatures extracted by the models. Scaled time loadings (ii, v) at time points were averaged per study group. Shaded area indicate standard errors of group means. Panels (iii, vi) present subject loadings compared across exposure groups using the Mann–Whitney U test with Bonferroni correction.

**S12 Fig. Replicability of subject-specific scaled time loadings in the COPSAC2010 study**. Panel (i) shows histograms of the average standard deviation, calculated for each subject by first finding the standard deviation of the scaled time loadings at each time point across submodels, and then taking the average. Panel (ii) shows the mean (lines) ±2 standard deviations (shaded areas) of scaled time loadings calculated at each time point across submodels for three representative subjects from the VD - no IAP group in the COPSAC_2010_ cohort. Rows correspond to model components 1-5.

**S13 Fig. Replicability of PARAFAC2-based subject-specific scaled time loadings in the FARMM study**. Panel (i) shows histograms of the average standard deviation, calculated for each subject by first finding the standard deviation of the scaled time loadings at each time point across submodels, and then taking the average. Panel (ii) shows the mean (lines) ±2 standard deviations (shaded areas) of scaled time loadings calculated at each time point across submodels for three representative subjects from the EEN study group in the FARMM study. Rows correspond to model components 1-3.

## Acknowledgments

We express our deepest gratitude to the children and families of the COPSAC_2010_ cohort. We thank Frederiek Wesel and Roel van der Ploeg for the insightful discussions.

## Data availability

Data and code underlying the simulations are available at https://github.com/blzserdos/parafac2_microbiome and archived at https://doi.org/10.5281/zenodo.17674126. Shotgun metagenomic sequence data from the FARMM dataset [15] were deposited under BioProject with accession code PRJNA675301. Processed data were obtained from [8]. Individual-level clinical data from the COPSAC_2010_ cohort are not publicly available to protect participant privacy, in accordance with the Danish Data Protection Act and European Regulation 2016/679 of the European Parliament and of the Council (GDPR) that prohibit distribution even in pseudo-anonymized form. Data can be made available under a joint research collaboration by contacting COPSAC’s data protection officer.

