## Supplementary Material for "Extracting host-specific developmental signatures from longitudinal microbiome data"

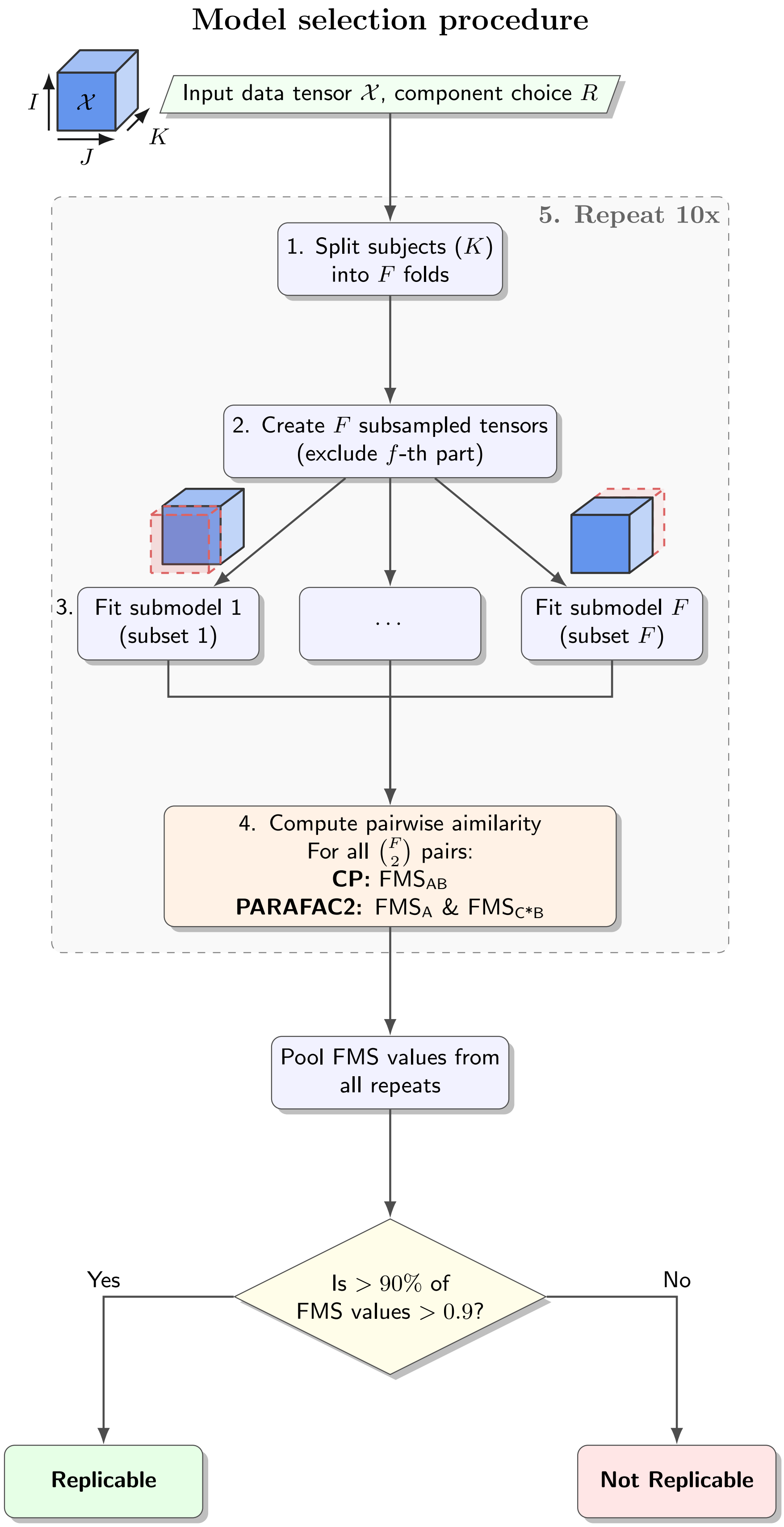


S1 Fig: Schematic of assessing the replicability of a CP or PARAFAC2 model with $R$ components used in model selection.

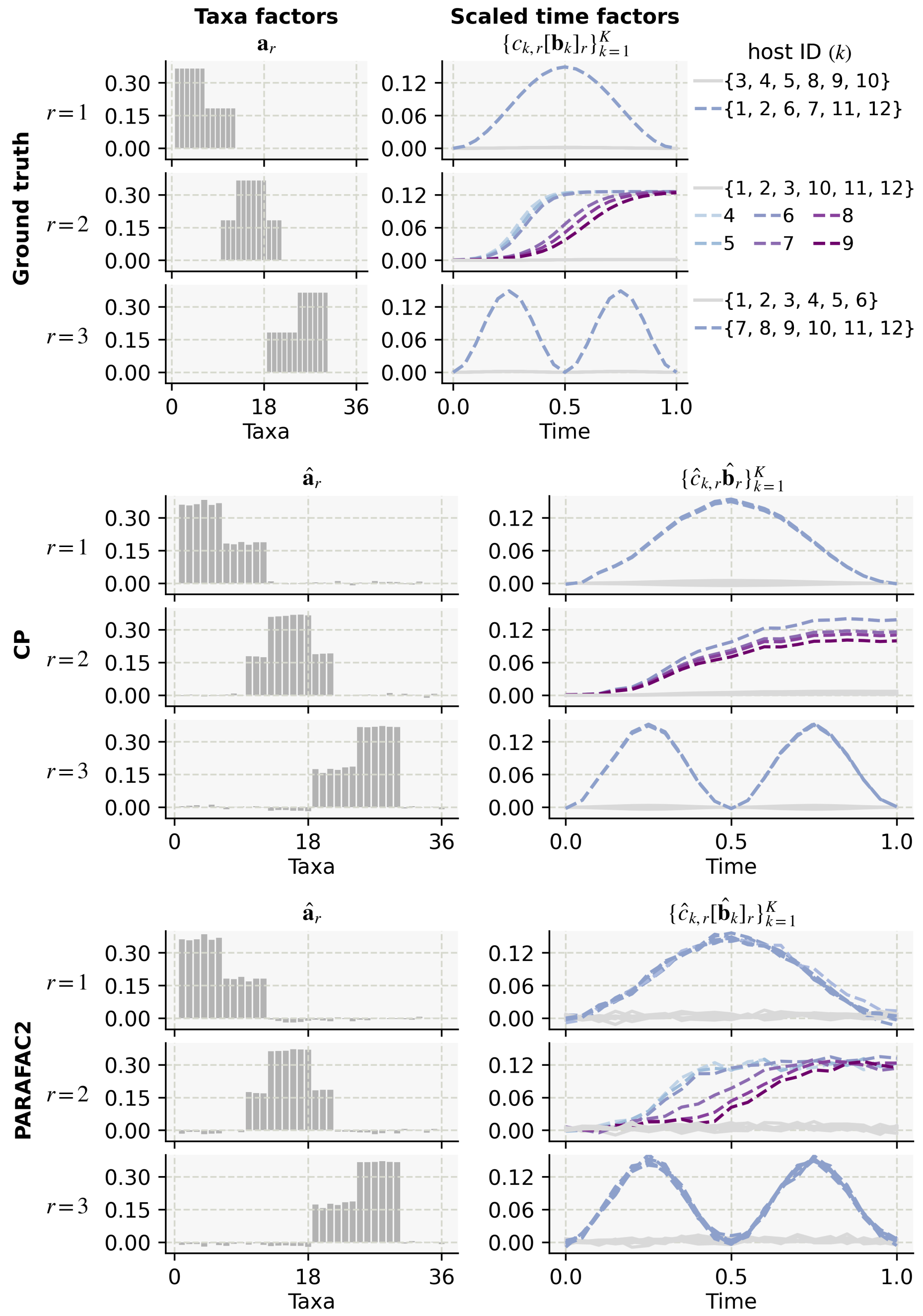


S2 Fig: Ground truth factors and their recovery by CP and PARAFAC2. Noise was added to the tensor as $\mathcal{X}_{noisy} = \mathcal{X} + \eta \|\mathcal{X}\| \frac{\mathcal{E}}{\|\mathcal{E}\|}$ with $\mathcal{E} \sim \mathcal{N}(0, 1)$ and $\eta = 0.25$ where $\mathcal{X}$ is the simulated tensor. Top panel: ground truth factors including the taxa ($\mathbf{a}_r$), time ($[\mathbf{b}_k]_r$), and subjects ($\mathbf{c}_r$) loadings with the time loadings scaled by the respective subject loading ($c_{k,r}[\mathbf{b}_k]_r$). The microbial signature $\mathbf{a}_r$ is present in subject $k$ over time according to their scaled time loadings $c_{k,r}[\mathbf{b}_k]_r$. The presence/absence of the signature in the subject is indicated by blue dashed/gray continuous line. The patterns in component 2 contain subject-specific time loadings $c_{k,2}[\mathbf{b}_k]_2$ for $k \in \{4, 5, 6, 7, 8, 9\}$ as highlighted by the respective line colors. Middle and bottom panels: factors recovered by CP and PARAFAC2, respectively.

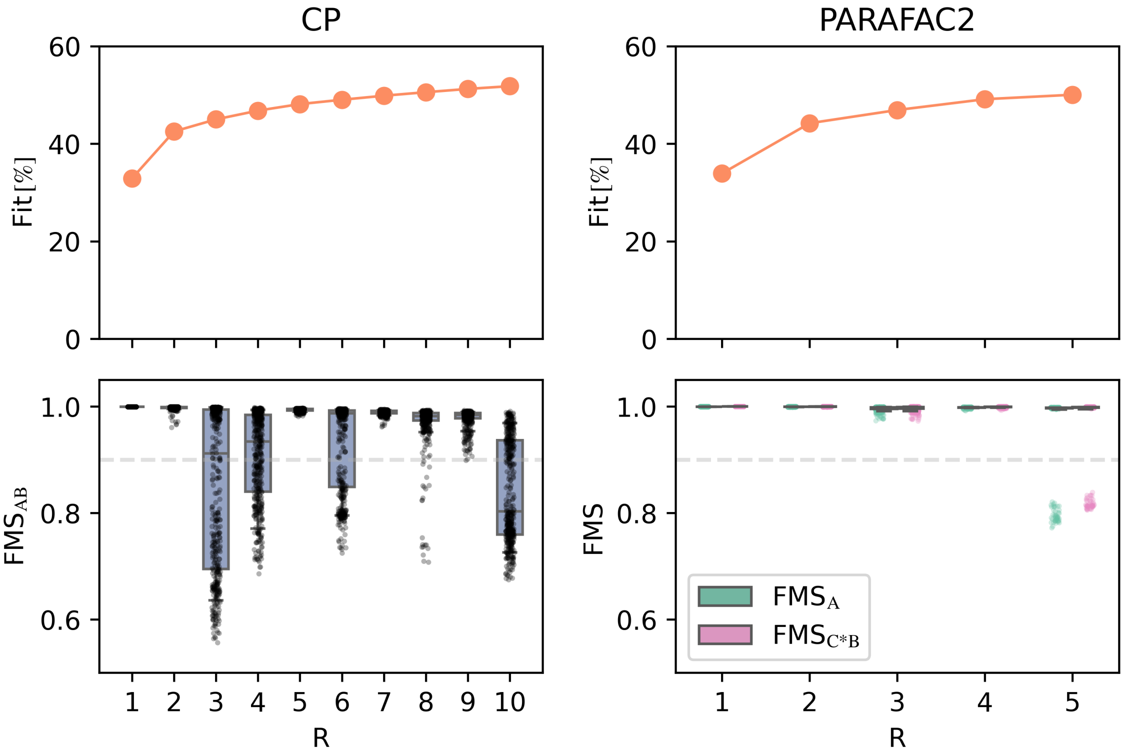


S3 Fig: Selection of the number of components for CP and PARAFAC2 models of $COPSAC_{2010}$ data. Model fit (top row) and replicability (bottom row) in CP and PARAFAC2 models with increasing number of components ($R$). Box plots represent the median and interquartile range, with whiskers extending to the 10th and 90th percentiles. FMS: factor match score. While a 9-component CP model also satisfied our replicability criteria, the components related to early life and our exposures of interest were similar between the models, therefore we selected the 7-component CP model for parsimony.

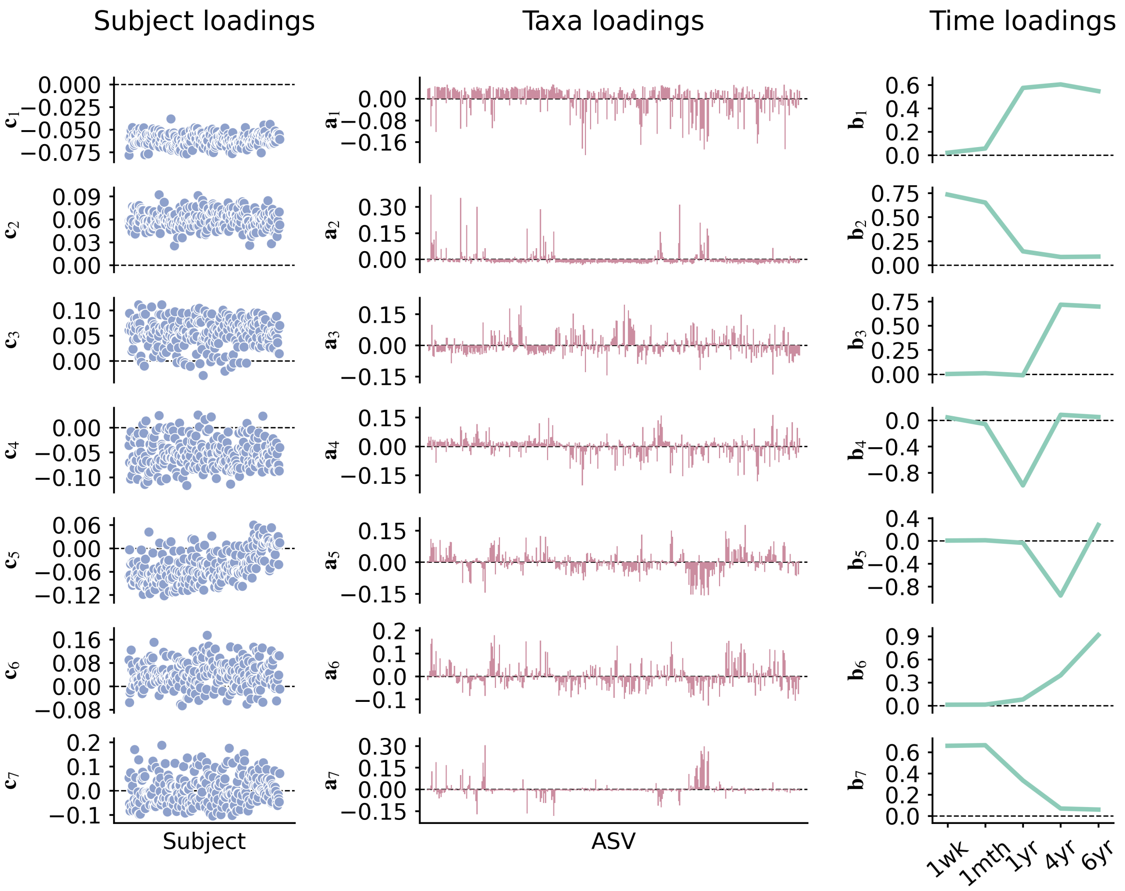


S4 Fig: Seven-component CP model of the COPSAC$_{2010}$ data. Each row shows the subject loadings ($\mathbf{c}_r$), taxa loadings ($\mathbf{a}_r$), and time loadings ($\mathbf{b}_r$) for the $r$th component.

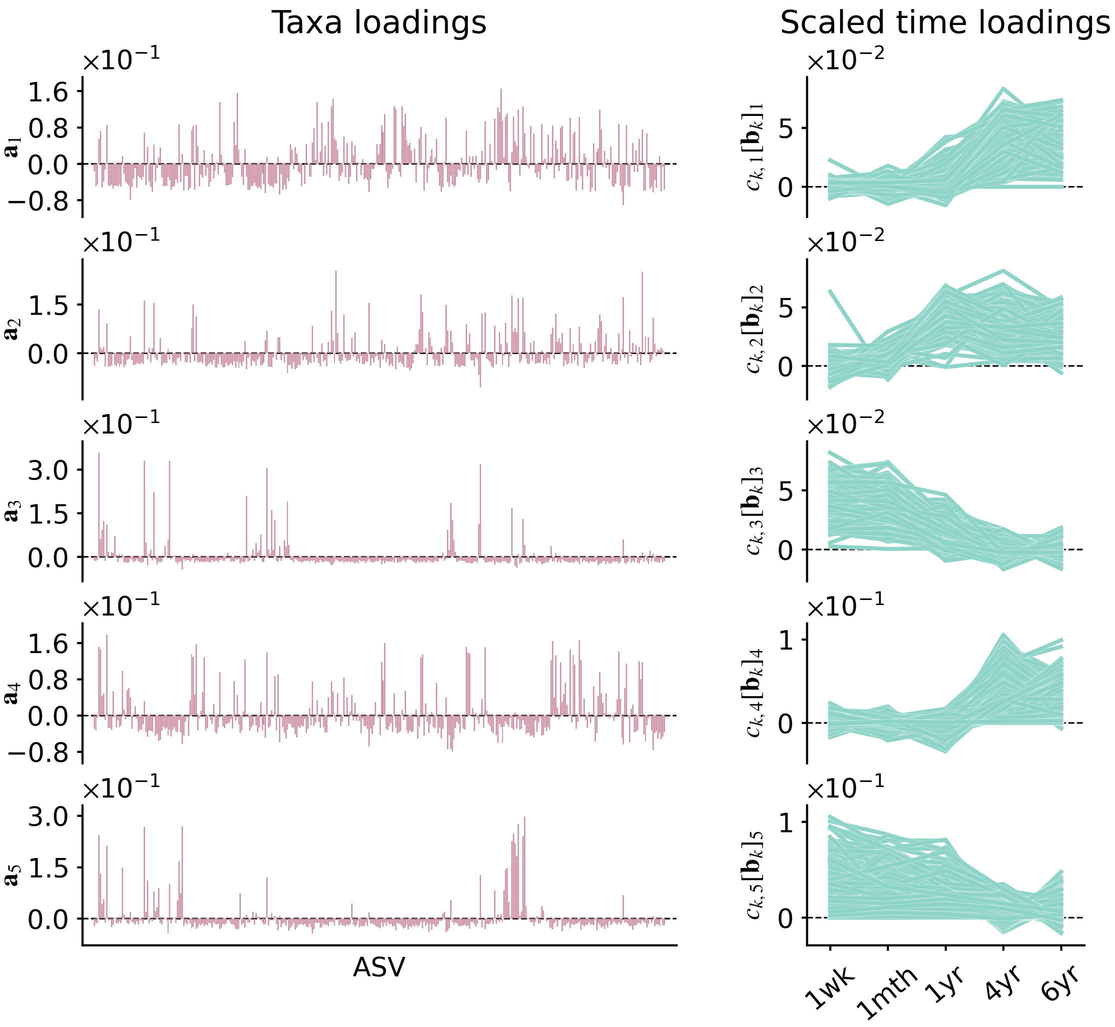


S5 Fig: Five-component PARAFAC2 model of the $\mathrm{COPSAC}_{2010}$ data. Each row shows a component comprising taxa loadings ($\mathbf{a}_r$) and the time loadings scaled by their corresponding subjects loadings ($c_{k,r}[\mathbf{b}_k]_r$) for the $k$th subject and $r$th component.

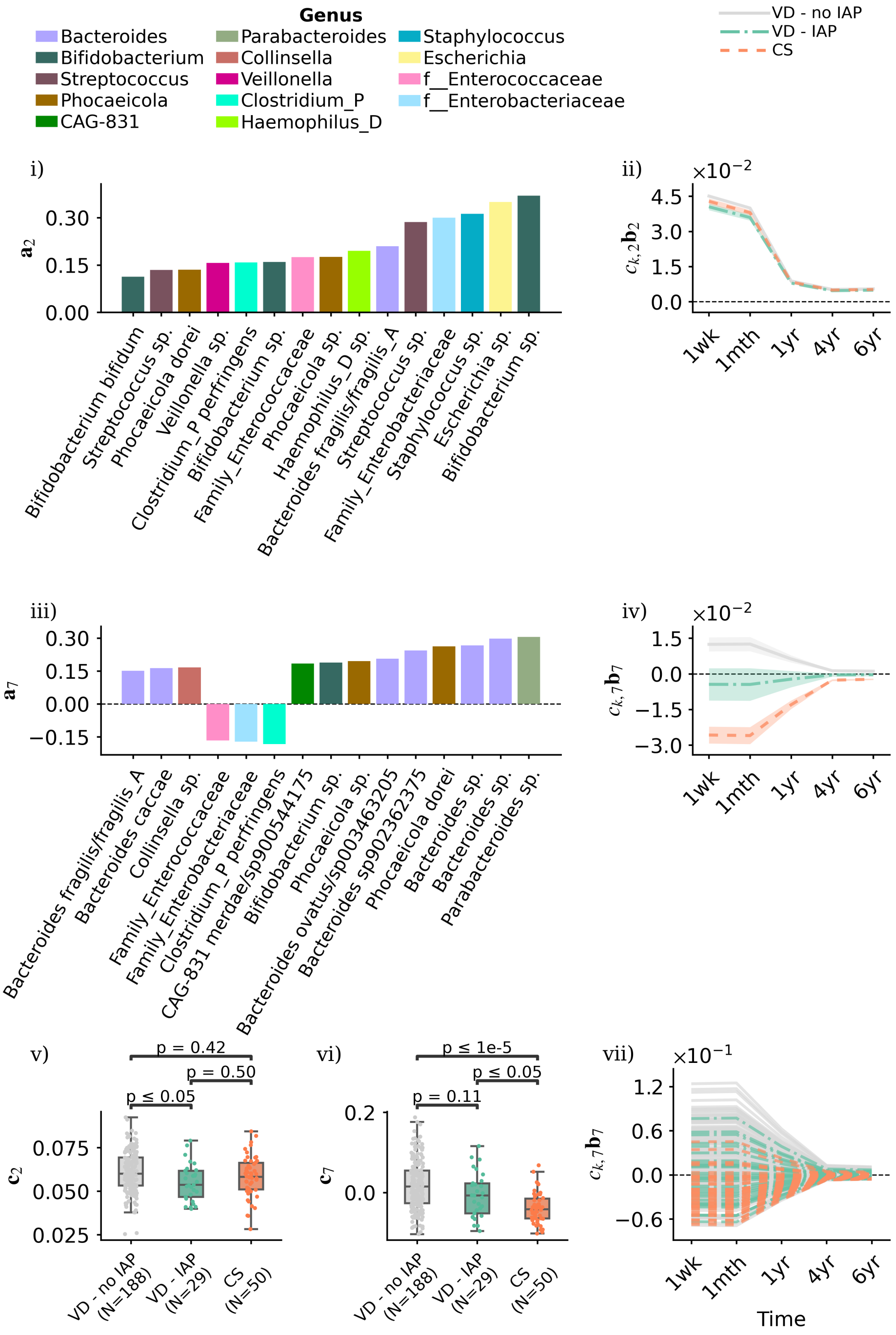


S6 Fig: Early life-related components of the selected CP model of the COPSAC$_{2010}$ cohort. Components 2 (i–ii) and 7 (iii–iv) capture shifts in gut microbiome composition during the first year of life. Panels (i, iii) show taxa loadings with the 15 ASVs of largest absolute weight in each component, colored by genus. Scaled time loadings (ii, iv) at time points were averaged per exposure group. Shaded area indicate standard errors of group means. Panels (v, vi) present subject loadings in components 2 and 7 compared across exposure groups using the Mann–Whitney U test with Bonferroni correction. Panel (vii) illustrates subject-specific scaled time loadings of component 7.

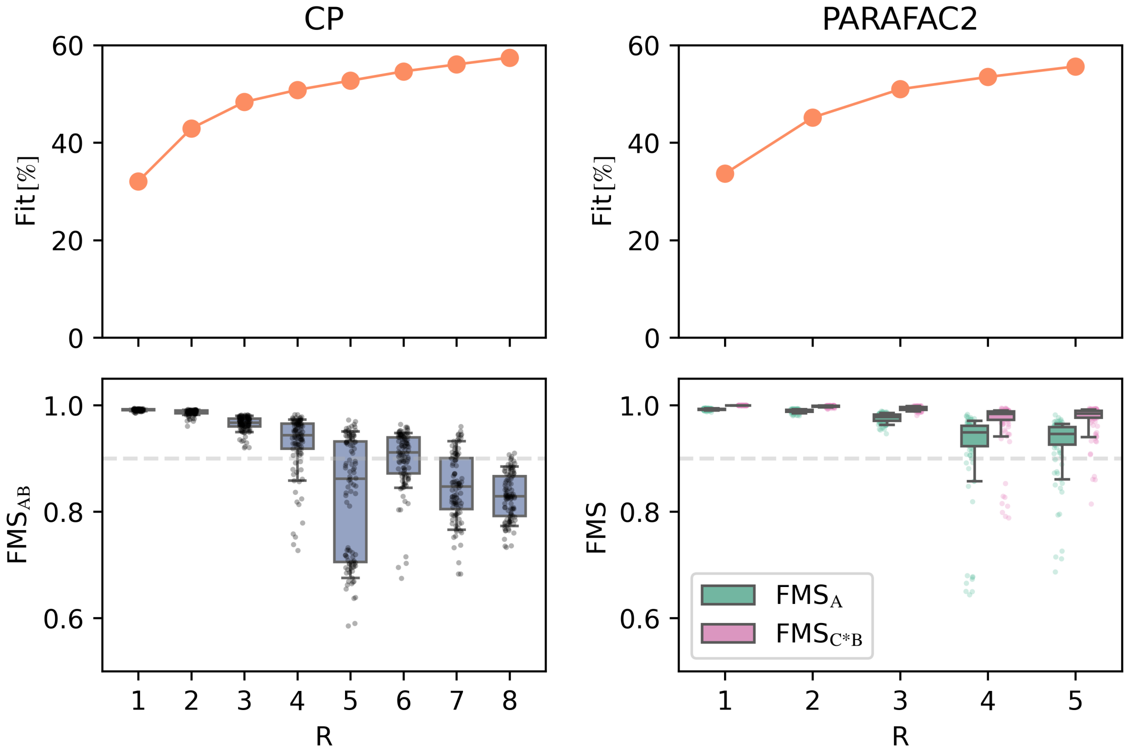


S7 Fig: Selection of the number of components for CP and PARAFAC2 models of FARMM study data. Model fit (top row) and replicability (bottom row) in CP and PARAFAC2 models with increasing number of components ($R$). Box plots represent the median and interquartile range, with whiskers extending to the 10th and 90th percentiles. FMS: factor match score.

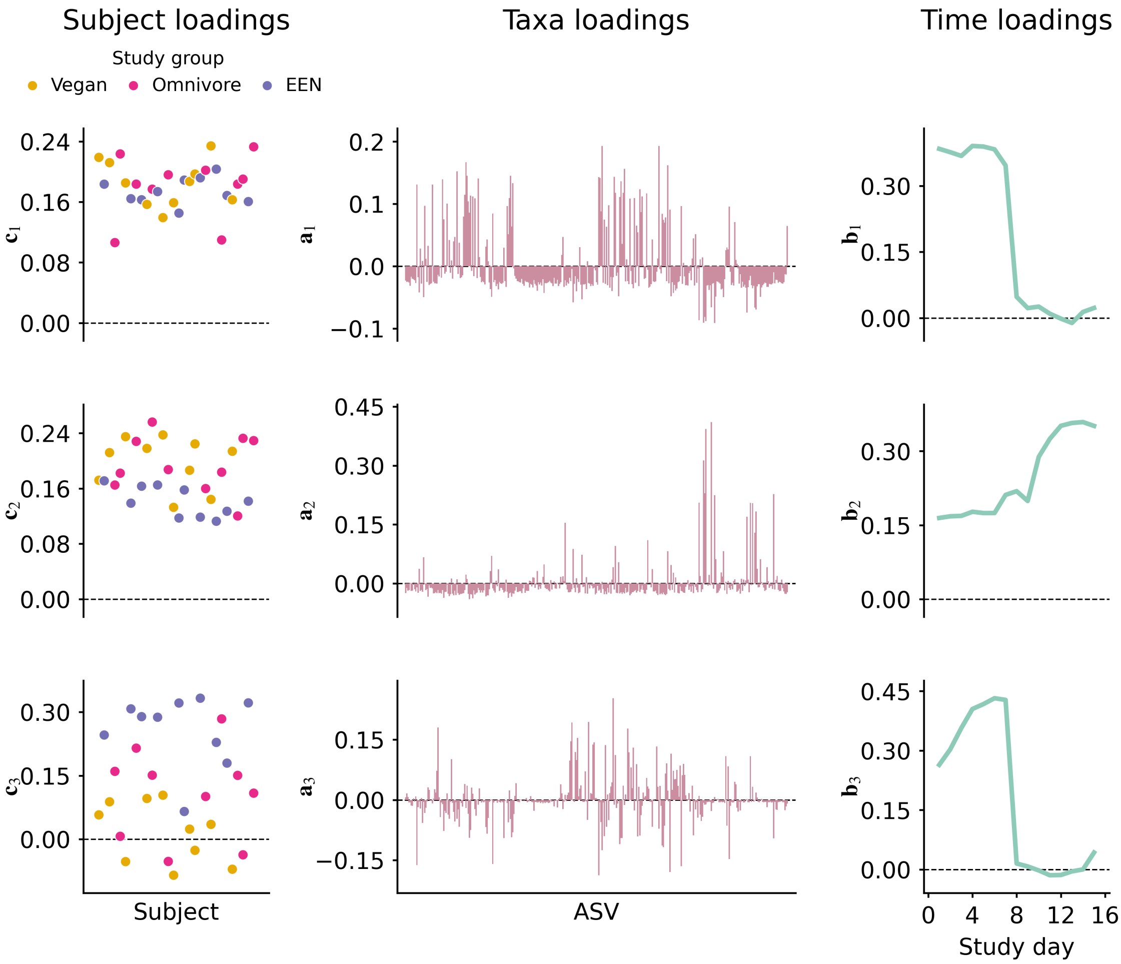


S8 Fig: Three-component CP model of the FARMM study data. Each row displays the subject loadings ($\mathbf{c}_r$), taxa loadings ($\mathbf{a}_r$), and time loadings ($\mathbf{b}_r$) for the $r$th component.

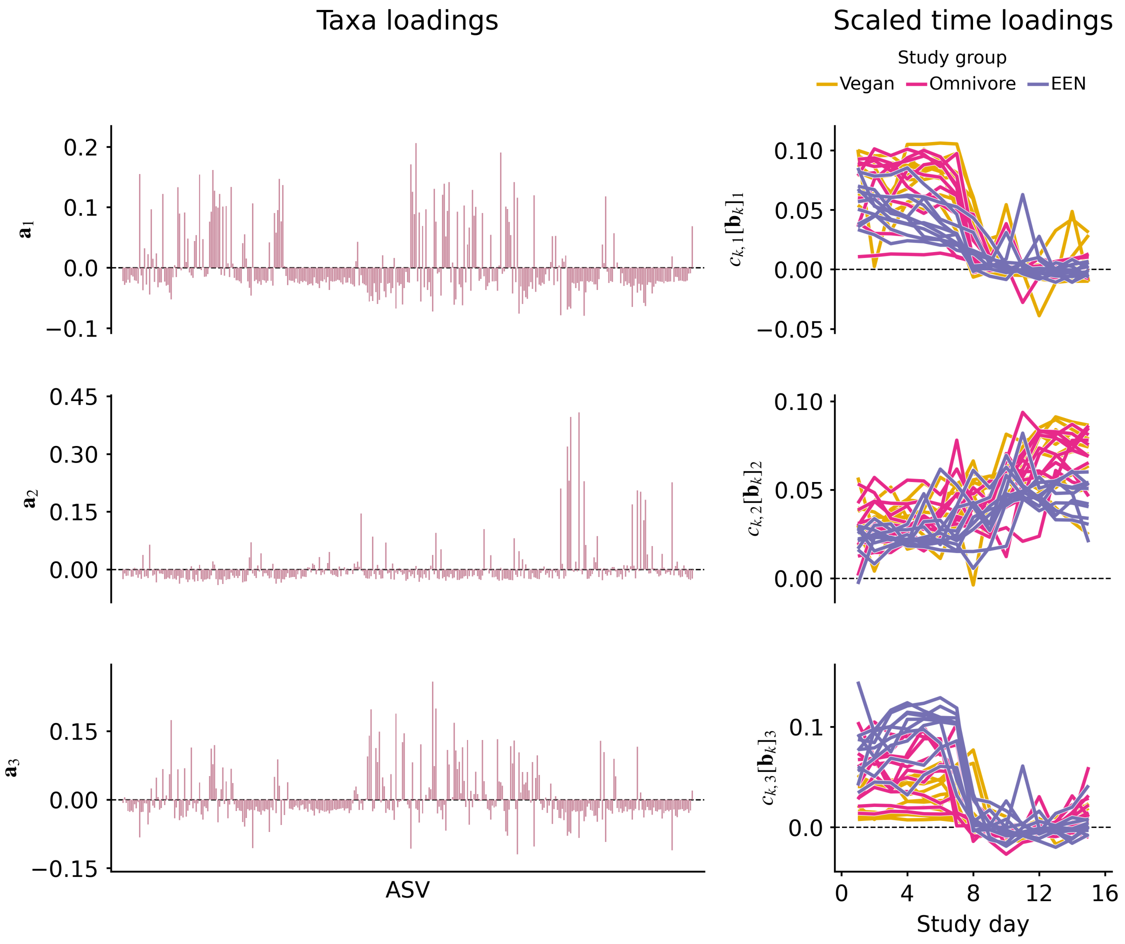


S9 Fig: Three-component PARAFAC2 model of the FARMM study data. Each row displays the taxa loadings ($\mathbf{a}_r$) together with the time loadings scaled by their corresponding subject loadings ($c_{k,r}[\mathbf{b}_k]_r$) for the $k$th subject and $r$th component.

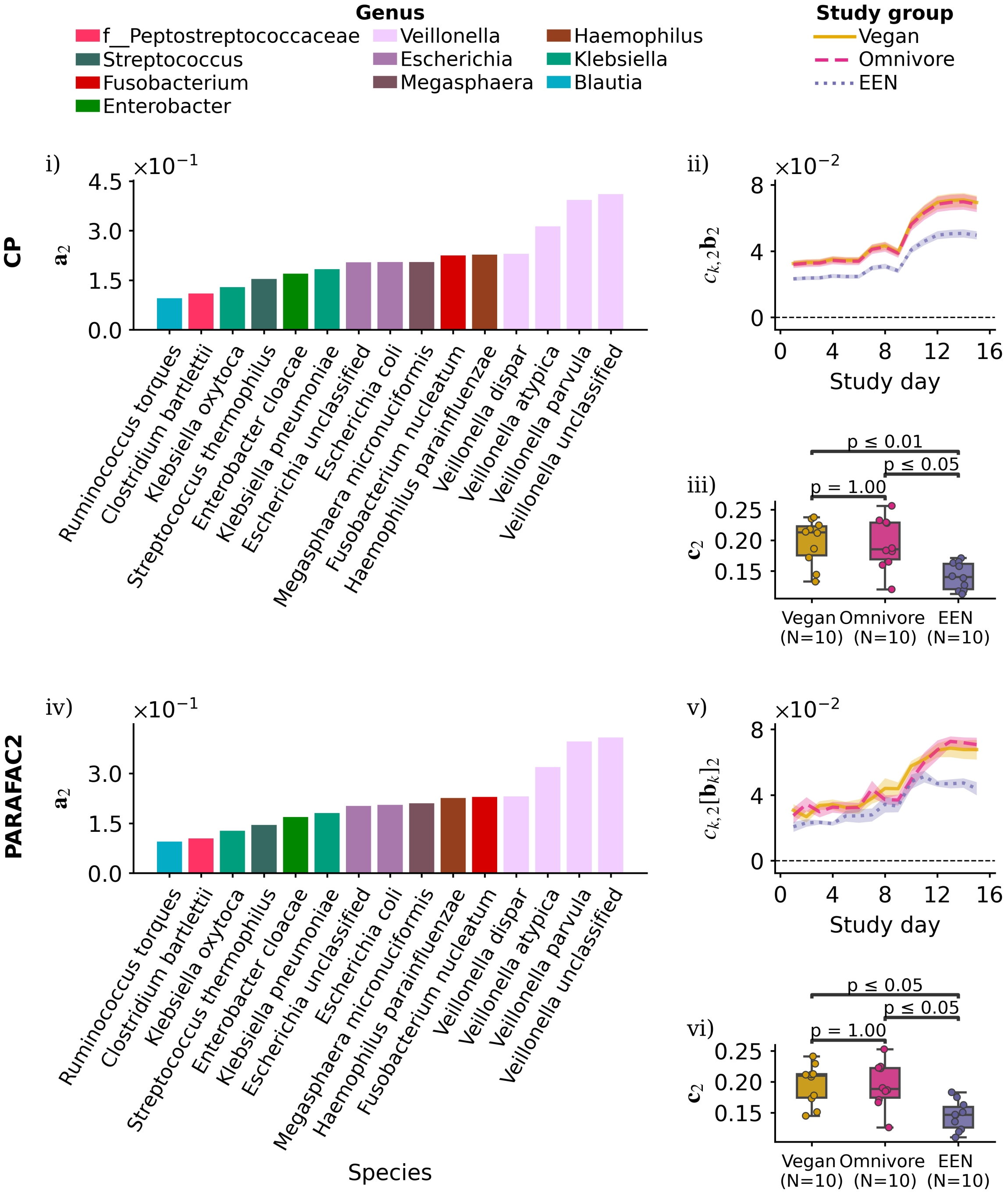


S10 Fig: Component 2 of the selected CP (panels i–iii) and PARAFAC2 (panels iv–vi) models, both associated with the recovery phase (days 9–15) of the trial. Panels (i, iv) display the 15 species with the largest absolute weights in each component, representing the microbial signatures extracted by the models. Scaled time loadings (ii, v) at time points were averaged per study group. Shaded area indicate standard errors of group means. Panels (iii, vi) present subject loadings compared across exposure groups using the Mann–Whitney U test with Bonferroni correction.

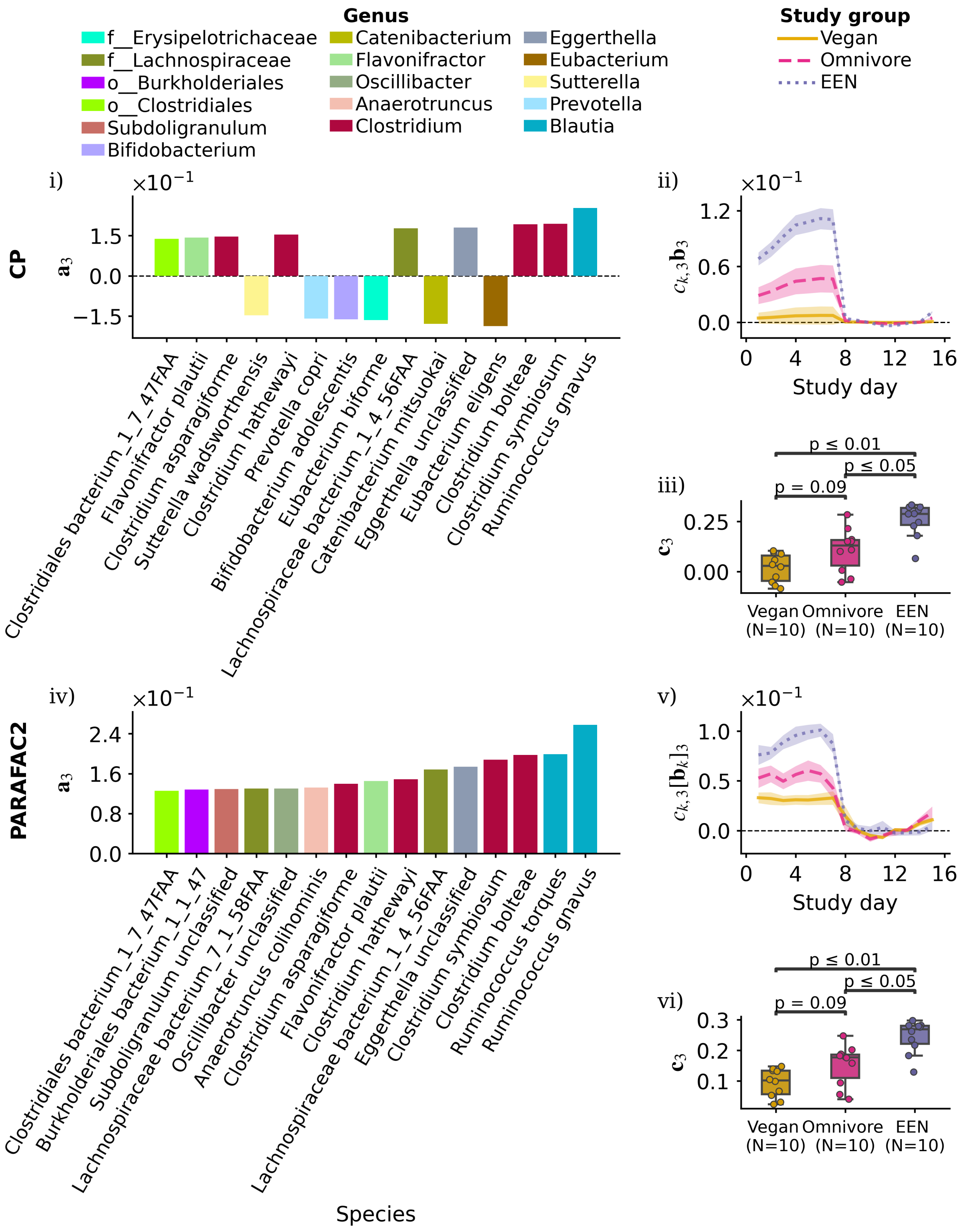


S11 Fig: Component 3 of the selected CP (panels i–iii) and PARAFAC2 (panels iv–vi) models, both associated with the dietary phase (days 1–5) and the antibiotics phase (days 6–8) of the trial. Panels (i, iv) display the 15 species with the largest absolute weights in each component, representing the microbial signatures extracted by the models. Scaled time loadings (ii, v) at time points were averaged per study group. Shaded area indicate standard errors of group means. Panels (iii, vi) present subject loadings compared across exposure groups using the Mann–Whitney U test with Bonferroni correction.

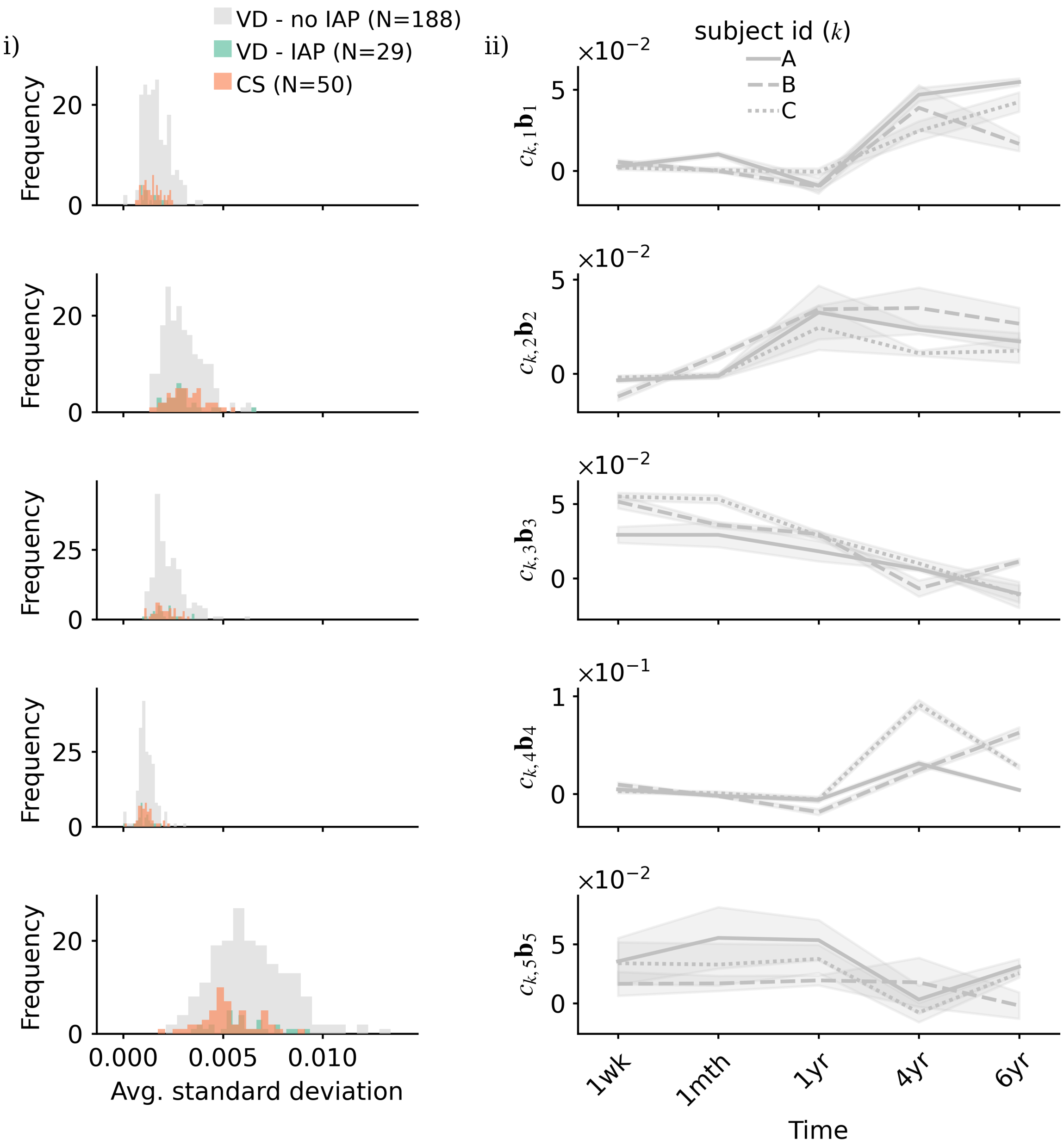


S12 Fig: Replicability of subject-specific scaled time loadings in the COPSAC2010 study. Panel (i) shows histograms of the average standard deviation, calculated for each subject by first finding the standard deviation of the scaled time loadings at each time point across submodels, and then taking the average. Panel (ii) shows the mean (lines) ±2 standard deviations (shaded areas) of scaled time loadings calculated at each time point across submodels for three representative subjects from the VD - no IAP group in the COPSAC$_{2010}$ cohort. Rows correspond to model components 1-5.

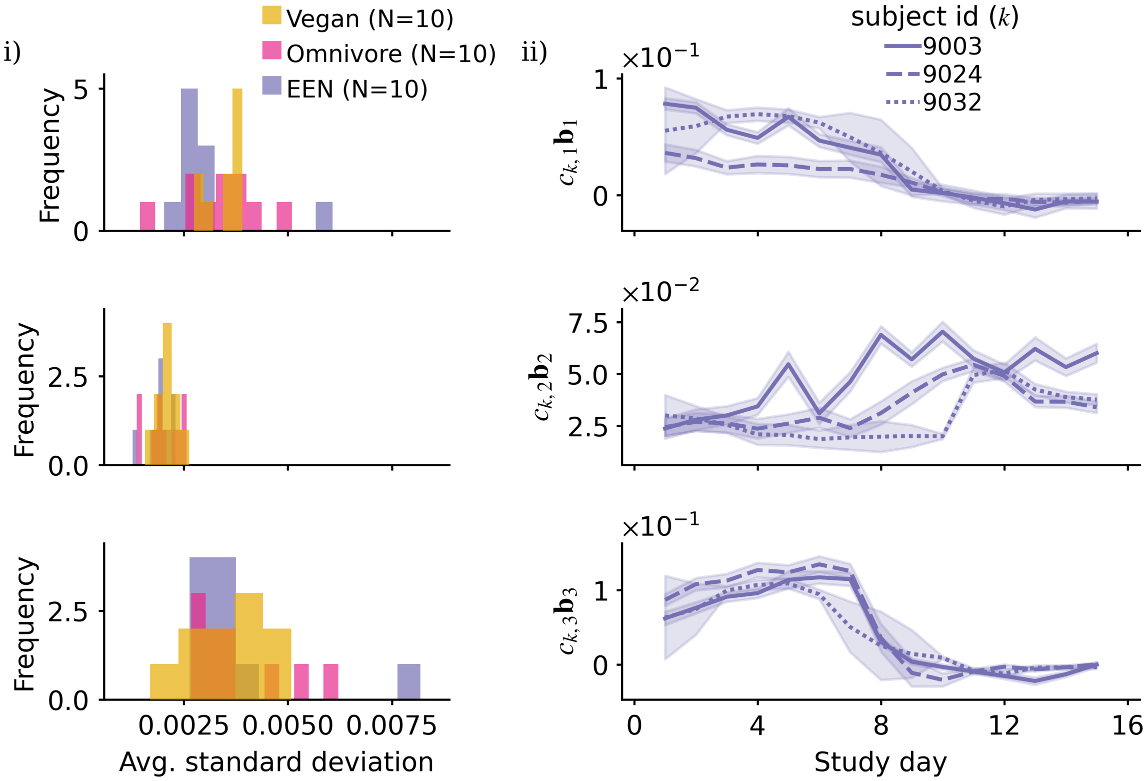


S13 Fig: Replicability of PARAFAC2-based subject-specific scaled time loadings in the FARMM study. Panel (i) shows histograms of the average standard deviation, calculated for each subject by first finding the standard deviation of the scaled time loadings at each time point across submodels, and then taking the average. Panel (ii) shows the mean (lines) $\pm 2$ standard deviations (shaded areas) of scaled time loadings calculated at each time point across submodels for three representative subjects from the EEN study group in the FARMM study. Rows correspond to model components 1-3.
